# Towards light responsive hydrogel-based valves for flow regulation

**DOI:** 10.1101/2025.11.06.686031

**Authors:** Annina Mittelholzer, Vincent Hickl, Katharina Maniura-Weber, Luciano F. Boesel, René M. Rossi, Markus Rottmar, Yashoda Chandorkar

## Abstract

Smart hydrogels are promising materials for soft actuators in biomedical applications thanks to their varied responses to external stimuli. Light is a particularly attractive trigger for contactless stimulation of hydrogels that can induce reversible morphological changes without damaging the fragile gels. To meet the varied needs of applications in microfluidics, soft robotics, biomedicine, and other fields, there is significant demand for novel valve designs that are highly tunable, miniaturizable, and respond quickly to stimuli while maintaining their function over many activation cycles. Additionally, it is crucial to develop a more quantitative understanding of the mechanics of valve operation in response to different stimuli, especially when active hydrogels are combined with other materials in multi-component devices. Here, stimuli-responsive valves are fabricated using active hydrogels deformed upon temperature changes and exposure to near-infrared radiation. Gold nanorods (AuNRs) acting as photothermal transducers are embedded inside N-isopropylacrylamide (NIPAM), allowing local morphological changes in response to light with high spatiotemporal control. These changes are described precisely as a function of the valve’s confinement, aspect ratio, and the parameters of the stimulus using quantitative image analysis, providing novel mechanistic insights. Changing the aspect ratio of the valves and the degree of confinement of the hydrogel causes valves to either open or close during heating and can be used to control the magnitude of their response to different stimuli. These varied morphological changes are due to local, inhomogeneous deformations of the gel. The use of light as a trigger enables local confinement of the valve, reversible opening and closing, and fast response times on the order of seconds. The valves are shown to withstand hydrostatic pressures of up to 18 kPa, providing high potential for biomedical applications where precise pressure control and quick switching between open and closed states is critical.

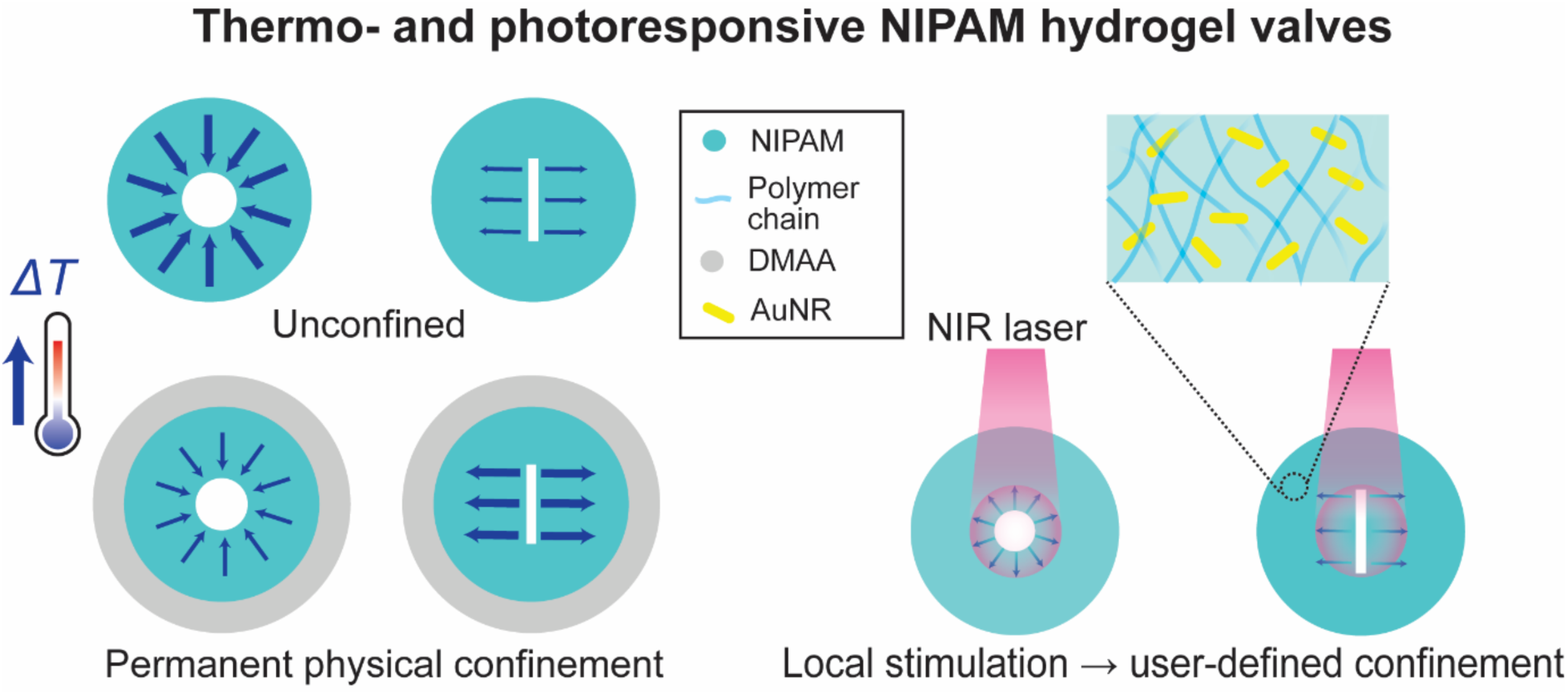

## INTRODUCTION

Hydrogels, known for their resemblance to human tissue, are promising materials for biomedical applications, including drug delivery, tissue engineering, and microfluidics for lab-on-chip technologies. The excellent compatibility of hydrogels with different biological entities means they can readily be incorporated with living tissue, while their physicochemical properties can be tuned through a variety of formulations and modifications for a wide range of applications [1], [2]. The plethora of existing hydrogel fabrication methods also provides extensive possibilities for their use in novel biomedical devices, where they can regulate cell development and behavior, control fluid flow, and achieve controlled drug release, to name but a few use cases [3]. Stimuli-responsive (or smart) hydrogels are hydrogel formulations that transform in response to changes in their environment such as temperature, pH, chemical composition, and pressure, or to a particular external signal such as mechanical stimulation, light, electromagnetic fields, sound etc. [4].

Stimuli-responsive (active) hydrogel valves are a particularly promising application for such active gels because, compared to passive valves, they allow for adjustable flow control and are less susceptible to irreversible clogging by particles [5]. Active hydrogel valves are growing in popularity, and different designs have been developed or proposed for a wide variety of applications, including microfluidics [6], soft robotics [7], drug delivery [8], and lab-on-chip technology [4]. A notable use case for active valves is implants for glaucoma patients, where they may be used to reduce intraocular pressure (IOP) via the drainage of excess fluid. Compared to conventional (passive) valves, an active valve can reduce the risk of hypotony, while the use of hydrogels provides improved biocompatibility compared to other implants [9], [10], [11]. Active hydrogel valves need to be highly responsive and reliable – opening and closing quickly, but only in response to a particular stimulus, even after multiple cycles [12]. They also need to be miniaturizable, both to allow effective incorporation into microfluidics, soft robots, or biomedical devices, and to reduce response times, which are dependent on the gel size. To meet the varied needs of different applications, it is highly advantageous if operation parameters such as the change in the valve outlet size, the response time, and the properties of the stimulus can be adjusted without fundamentally changing the valve design.

To meet these criteria, a major obstacle for the use of stimuli-responsive gels is their softness and resulting fragility, which makes the design of devices for consistent, long-term operation difficult. For the construction of hydrogel valves, reversible opening and closing without disintegration remains a challenge, especially when contact-based stimulation is used [12], [13]. One solution to these challenges is the use of pH sensitive valves [14], but this approach necessitates that all components of a multi component system are compatible with pH changes and hence restricts the usage of such systems. Temperature is a widely used trigger for thermoresponsive gels such as N-isopropylacrylamide (NIPAM), a very well researched polymer that undergoes a coil to globule transition with increase in temperature [15], [16], [17]. This transition is thermodynamically favored because the increase in temperature disrupts hydrogen bonds between the solvent and polymer moieties, thereby minimizing the total free energy. The volume phase transition temperature (VPTT) of NIPAM, where macroscopic gels shrink significantly, is typically around 32-34 °C.

Light is an attractive stimulus as it permits non-contact manipulation of hydrogels, allowing user-defined spatiotemporal control, and can induce reversible morphological changes without damaging fragile hydrogels [18], [19], [20]. Thermoresponsive gels like NIPAM can be made photoresponsive by combining them with a photothermal transducer: a component that heats up when irradiated by certain wavelengths of light. Previous studies have shown that gold nanorods (AuNRs) embedded in NIPAM will cause a local rise in temperature when part of the gel is illuminated by near infrared (NIR) light [21], [22]. A notable design for an active NIPAM valve is bio-inspired by plant stomata [23]: valves found on the lower epidermis of leaves of aerial plants, which open and close to regulate water and CO_2_. Like the guard cells which exert pressure on and modulate the behavior of plant stomata, a second gel is used to confine the NIPAM and thus regulate the opening and closing of the valve [24]. A relatively large (∼1 cm) version of this valve was shown to be operable with temperature, pH, or UV light stimuli, and was effective in preventing and allowing fluid to pass through below and above the VPTT, respectively [24]. Before such designs can be adapted to a particular application, the mechanics of hydrogel valves’ responses to different stimuli must be better understood to predict how changes to the valve shape, confinement, composition, and the properties of the stimulus will affect the valve’s before over multiple cycles. So far, the opening and closing of such valves have not been quantified as a function of temperature or hydrostatic pressure, meaning the underlying mechanics of their operation, and those of active hydrogel valves in general, remain poorly understood. Additionally, the use of UV stimulation may limit the design’s translational potential due to the harmful effects of UV radiation, with NIR light providing a promising alternative [25]. The size and response times (on the order of several minutes) must also be reduced significantly to make it suitable for time-sensitive applications.

Here, using a hydrogel valve design inspired by plant stomata, we precisely quantify how confinement of an active hydrogel by another, passive gel affects the valve’s operation. AuNRs are embedded inside NIPAM gels to act as photothermal transducers, enabling user control of the valve with both thermal and light stimuli. The extent of the valve’s opening and closing in response to both stimuli is described using quantitative image analysis as a function of the active gel’s confinement and the valve’s aspect ratio. The photoresponsive behavior of the valve is also quantified as a function of the parameters of the NIR illumination, demonstrating that this design provides excellent tunability and user control, as well as highly responsive and repeatable valve behavior. The valve’s resistance to hydrostatic pressure is tested to determine its suitability for biomedical applications, in particular for use in glaucoma implants. The comprehensive description of this valve design provides novel insights into the mechanics of hydrogel deformations under confinement and demonstrates its suitability for versatile uses.

**Figure 1.**
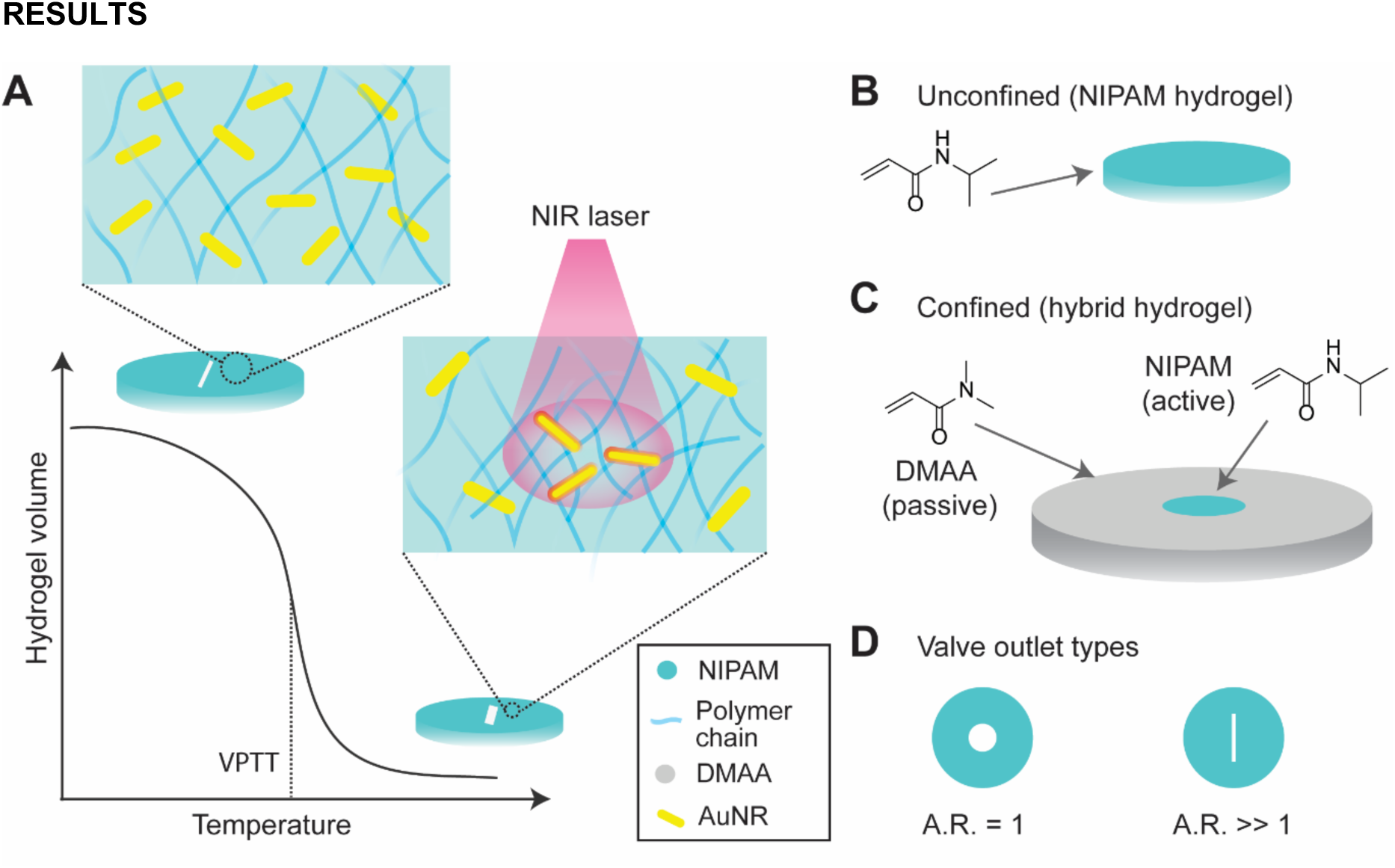
(A) Schematic showing the mechanism underlying the valve design. Light responsive hydrogel valves are fabricated from a thermoresponsive hydrogel (NIPAM) with embedded AuNRs. Owing to the photothermal properties of AuNRs, the valve outlet responds to NIR, causing the density of polymer chains to increase upon laser irradiation, leading to reversible opening and closing. (B) The simplest valve design is comprised of an unconfined NIPAM valve and (C) a confined NIPAM valve covalently bound to a passive shell of N,N’Dimethylacrylamide (DMAA). (D) Two different types of valve outlets were studied, with aspect ratio (A.R). = 1 i.e. circular outlets and A.R. >> 1, i.e. slit-shaped outlets. (Illustrations not to scale)

## RESULTS

Photo- and thermoresponsive hydrogel valves are created by cutting apertures inside a disc-shaped crosslinked NIPAM hydrogel into which AuNRs are embedded, as shown in Fig 1A. The volume of the NIPAM hydrogel decreases sharply when it is heated past the volume phase transition temperature (VPTT ≈ 34°C). Upon exposure to NIR light, the embedded AuNRs are rapidly heated to induce a volume phase transition. To test how confinement of the gel affects the operation of these valves, the active NIPAM hydrogel disc can be surrounded by an outer ring of a passive hydrogel – DMAA, the volume of which does not change significantly in response to temperature, regardless of whether an outlet is present or not (Fig S1). The confined valves are fabricated in such a way that the DMAA and NIPAM are crosslinked together at the interface (see Methods), ensuring the two gels cannot be separated. Valves are either circular (aspect ratio: *AR* = 1) or slit-shaped (*AR* > 10). This approach provides an ideal framework to study the varied responses of an active hydrogel valve to light exposure, changes in temperature, confinement, and morphology.

**Figure 2.**
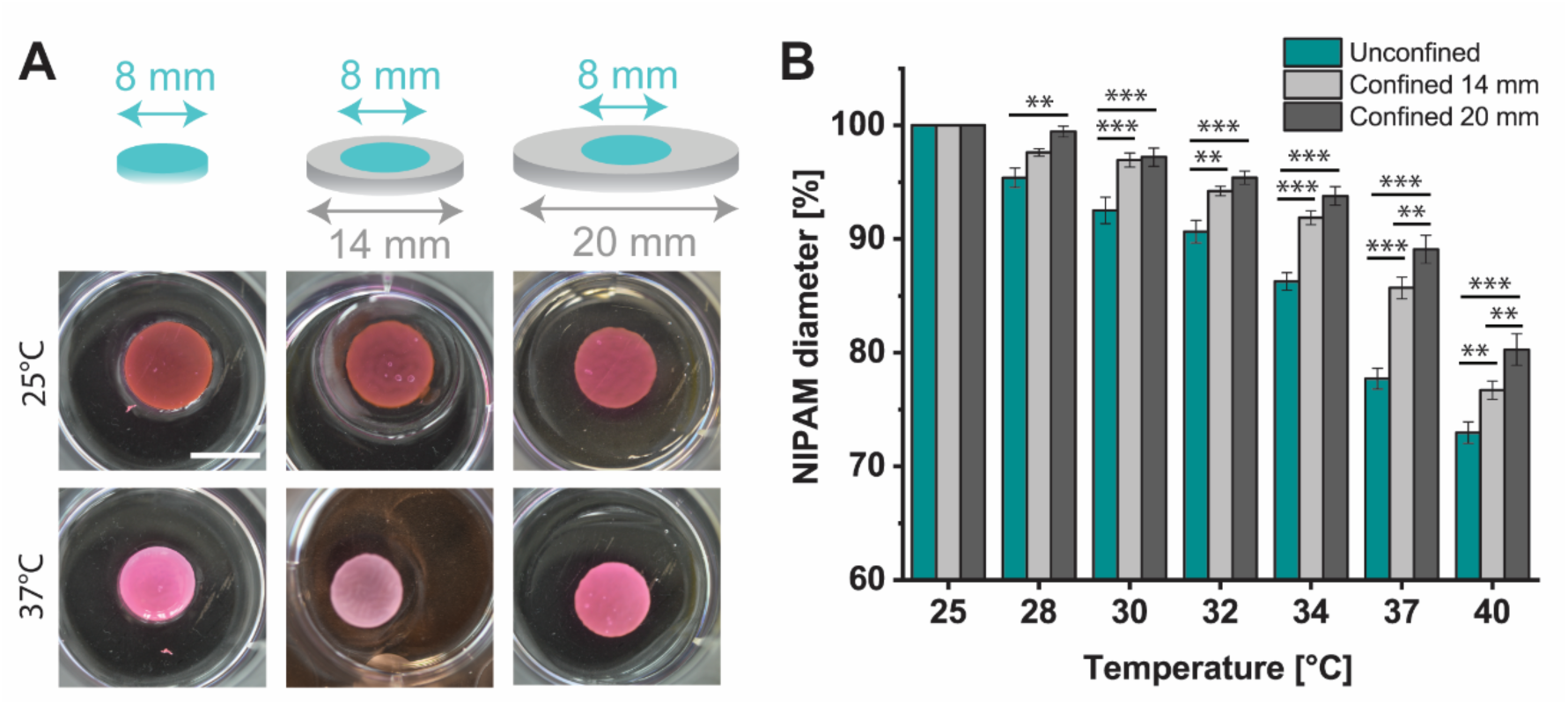
Effect of confinement on hydrogel response to heating. (A) Schematic showing the unconfined and confined bulk NIPAM gels (without outlet) with 2 different degrees of confinement (14 mm and 20 mm). Representative photos of gels (without slits) at 25 and 37 °C are shown. (B) Changes in the diameter of NIPAM gels in an unconfined and a confined state in response to changes in temperature, normalized to the diameter of the respective unconfined or confined NIPAM hydrogel at 25°C. Scale bar 1 cm. Statistical significance levels *, **, *** were determined using two-way ANOVA (factors of temperature and confinement) followed by Tukey multiple comparisons tests and represent p < 0.05, 0.01 and 0.001, respectively. Details of the statistical analysis are shown in Table S2.The data were collected from 6 independent samples for each condition.

The swelling of the thermoresponsive hydrogel in response to temperature can be controlled by confining it with a passive hydrogel. Disks of active hydrogel (either unconfined or confined by a ring of passive DMAA with a width of 3 mm or 6 mm, corresponding to a total diameter of 14 mm and 20 mm, respectively) were heated to various temperatures between 25 °C and 40 °C (Fig 2A). In all cases, the diameter of the active NIPAM decreases monotonically with increasing temperature (Fig 2B). The effect is non-linear, with the diameter decreasing more sharply above 34 °C, and well-described by a logistic function (*R*^2^ ≥ 0.98, see Fig S2 and Table S1). The diameter of the unconfined gel becomes significantly smaller than the gels confined by 14 mm and 20 mm of DMAA for all temperatures above 28 °C and 30°C, respectively. At 37 °C and 40 °C, both the unconfined and the confined 14 mm gels were significantly smaller than the confined 20 mm gel. This result shows that confinement of the active gel by a passive ring alters its morphological response to a change in temperature, which may be used to control the operation of a hydrogel valve. When confined, the contraction of the active gel at the VPTT is inhibited by the physical link to the passive gel, which is due to the covalent bonds formed during cross-linking.

**Figure 3.**
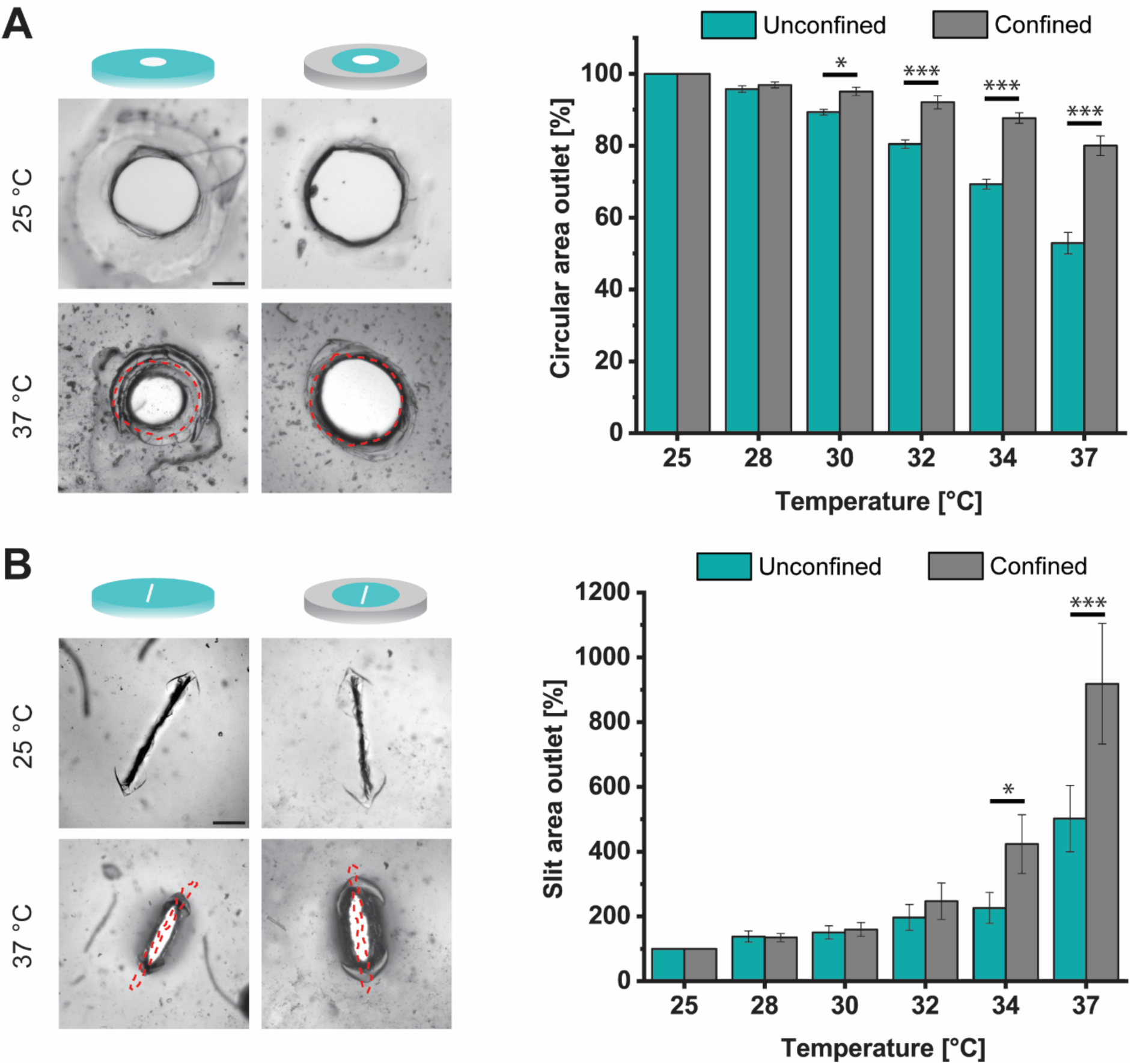
Valve outlet area as a function of temperature, confinement, and outlet aspect ratio. (A) NIPAM valves with circular outlet, (B) NIPAM valves with slit-shaped outlet. Red dashed lines represent the valve outlet outline at 25 °C. Illustrations not to scale, scale bar 200 µm. Error bars represent standards errors; *, **, *** were determined using two-way ANOVA (factors of temperature and confinement) followed by Tukey multiple comparisons tests and represent statistical significance at p < 0.05, 0.01 and 0.001, respectively. Details of the statistical analysis are shown in Tables S3 and S4. Scale bar 200 µm. The data were collected from 3 experiments with 2 or 3 replicates each (8 or 9 independent samples for each condition).

The aspect ratio of the thermoresponsive hydrogel valve determines whether the size of the opening increases or decreases with rising temperature. Hydrogels with circular and slit-shaped valve outlets, with and without a confining ring of DMAA, were heated from 25 °C to 37 °C, as before. The circular valve outlet area decreases with rising temperature. At 37 °C, the area reaches 53 ± 2% and 80 ± 1% of its value at 25 °C for the unconfined and confined case, respectively (Fig 3A). On the contrary, the area of the slit-shaped outlet valve increases (Fig 3B). At 37 °C, the area of the slit increases by 502 ± 89% and 897 ± 134% for the unconfined and confined case, respectively. Across all replicates, the largest increase observed was about 1200%. Thus, changing the aspect ratio of a hydrogel valve fundamentally alters its response to changes in temperature, and can be used to control whether the valve opens or closes when heated.

Confinement of the thermoresponsive hydrogel valve by a passive hydrogel can be used to alter its response to changes in temperature. For gels with a circular opening, the decrease in the area in response to temperature is significantly reduced when the NIPAM is confined with a ring of DMAA (Fig 3A). On average, the circular outlets in the unconfined gels shrink more than twice as much as those in confined gels compared to their respective sizes at 25 °C. The difference is significant for all measured temperatures ≥ 30 °C. On the other hand, the increase in area of the slit-shaped outlets is greater for the gels confined with a ring of DMAA (Fig 3B). In this case, the difference is significant for ≥ 34 °C, above the VPTT. For both aspect ratios, confinement of the active gel results in larger outlet areas above the VPTT compared to the unconfined case (the circular valve closes to a lesser extent, and the slit valve opens further). Performing two-factor ANOVA reveals that for both valve shapes, the effects of temperature, confinement, and their interaction all have significant effects on the outlet area (Table S4).

These results show that the operation of an active hydrogel valve depends on the complex interplay between its aspect ratio and the way it is confined by its surroundings. The aspect ratio of the valve determines whether it opens or closes in response to an increase in temperature, while the degree of confinement of the gel determines the extent of the change in outlet area. Considering these effects is especially important when incorporating a valve into a multi component device, where the active hydrogel will necessarily be confined by other components.

**Figure 4.**
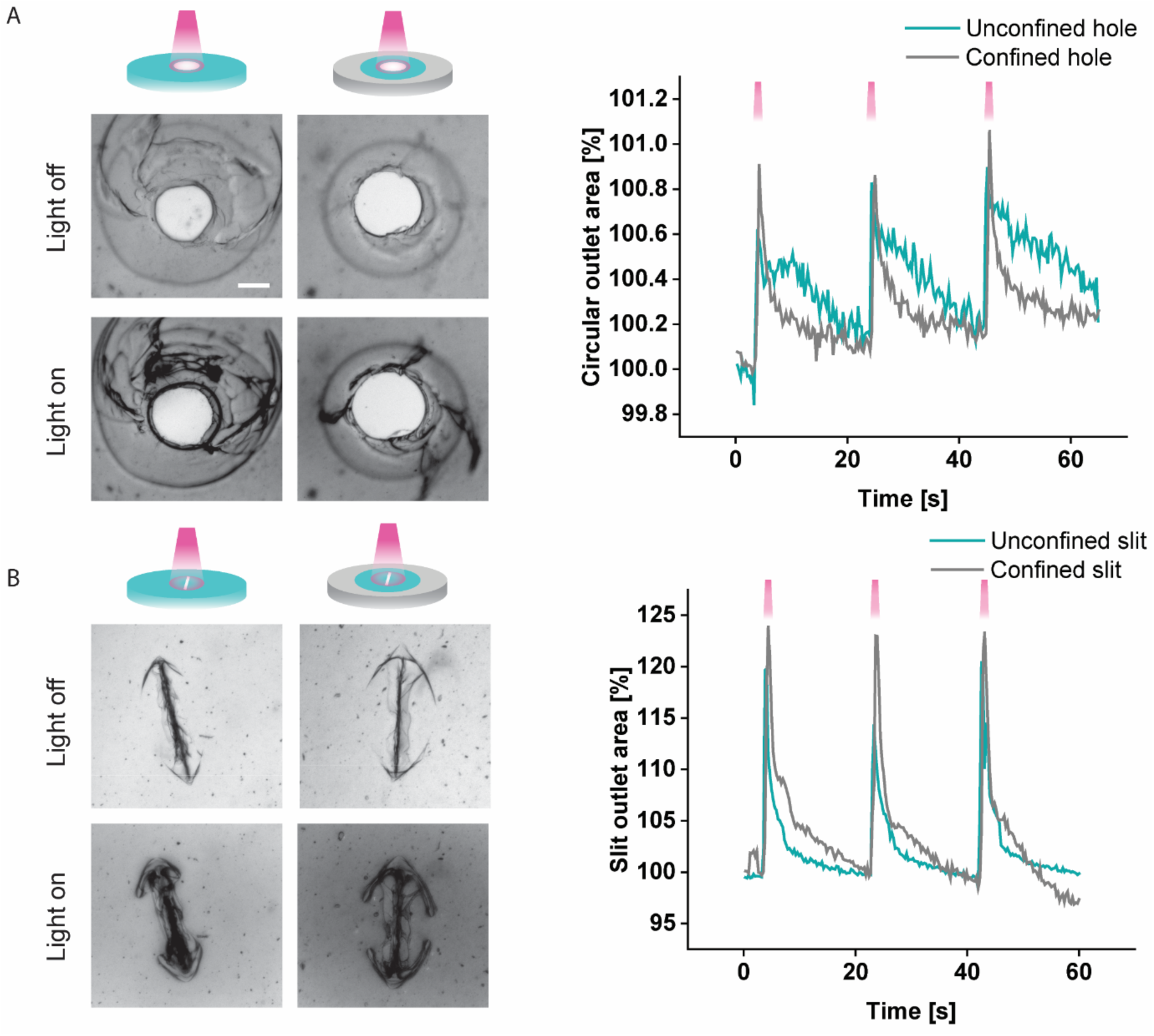
Valve response to NIR light. (A) Left: Images of the circular outlets in confined and unconfined valves without (top) and with (bottom) irradiation (800 nm, 152 mW, on time 500 ms, 0.05 Hz, interval 20 s). Right: Valve opening area as a percentage of its initial (non-illuminated) value for 3 consecutive illumination pulses. (B) Left: Images of the unconfined and confined slit outlet under the same conditions as (A). Right: Slit valve area as a percentage of its initial value for three illumination pulses. Scale bar 200 µm. Each valve was constructed and tested in triplicate.

Both circular and slit-shaped valve outlets are enlarged upon exposure to NIR light. To test the photoresponsive behavior of the NIPAM valve containing AuNRs, a NIR laser was used to illuminate the area immediately surrounding the outlet (Fig 4 and Fig S3), causing only that region of the gel to be heated, as confirmed by a NIR temperature camera (Fig S4). The circular region irradiated by the laser has a diameter of 1.4 mm. Unlike when the entire gel is heated, the outlet increases in size regardless of its aspect ratio and whether the NIPAM gel is confined with a ring of DMAA (Fig 4A). For the circular hole, the area increases by only 0.8 ± 0.1% in the unconfined case and 0.9 ± 0.1% in the confined case, whereas for the slit, the area increases by 20.3 ± 0.1% and 23.5 ± 0.3% in the unconfined and confined cases, respectively (Fig 4B). Here, as with the case where the whole gel is heated, the slit area increases more in the confined case. While all valves constructed here respond to the light stimulus, the effect observed is much stronger for the slit, further highlighting the importance of the valve’s aspect ratio. The opening and closing of these valves were consistent across 3 independent replicates for each condition and multiple illumination cycles (Fig S5). The opening of a confined slit valve is shown in supporting video 1. Future work to adapt this design for specific applications requires testing the valve over a much larger number of cycles (hundreds to thousands), under conditions relevant to the particular use case.

The difference between temperature and light as a trigger may be explained by the localized exposure to light: since only the part of the hydrogel being irradiated is heated, the surrounding hydrogel essentially functions as a passive hydrogel and provides physical confinement to the outlet. For a circular opening, this high degree of confinement causes the valve to increase in size because the central, illuminated portion of the hydrogel shrinks, while the volume of the unheated portion remains constant. Thus, the use of NIR illumination provides greater spatial control of the valve, making it possible to achieve characteristics of physical confinement upon demand without the need for passive hydrogel elements, thereby facilitating miniaturization of these devices for biomedical applications.

After exposure to NIR light, both types of valve outlets return to their original size and can be reopened upon repeated illumination. For all valves constructed here – circular and slit-shaped outlets with and without confinement – the valve opens sharply within less than 1 s of the laser being turned on and closes quickly after the NIR light is turned off. Within 20 s, the area of the outlet returns approximately to its original size. For the circular hole, the area returns to within 0.2% of its original size (Fig 4A). This value is a fifth of the maximum opening, suggesting slightly more time may be required between cycles. For the slit, the valve closes completely (the area returns to 100%) after each cycle (Fig 4B). Then, when the light is turned on again, the valve rapidly opens again. No qualitative difference in the opening or closing of the valve is observed over subsequent illumination cycles with NIR light, showing that this design provides highly reliable valve operation, regardless of aspect ratio and type of confinement. The slit valve had a significantly stronger response to NIR illumination, and, because of its high aspect ratio, is better suited to prevent flow when closed.

**Figure 5.**
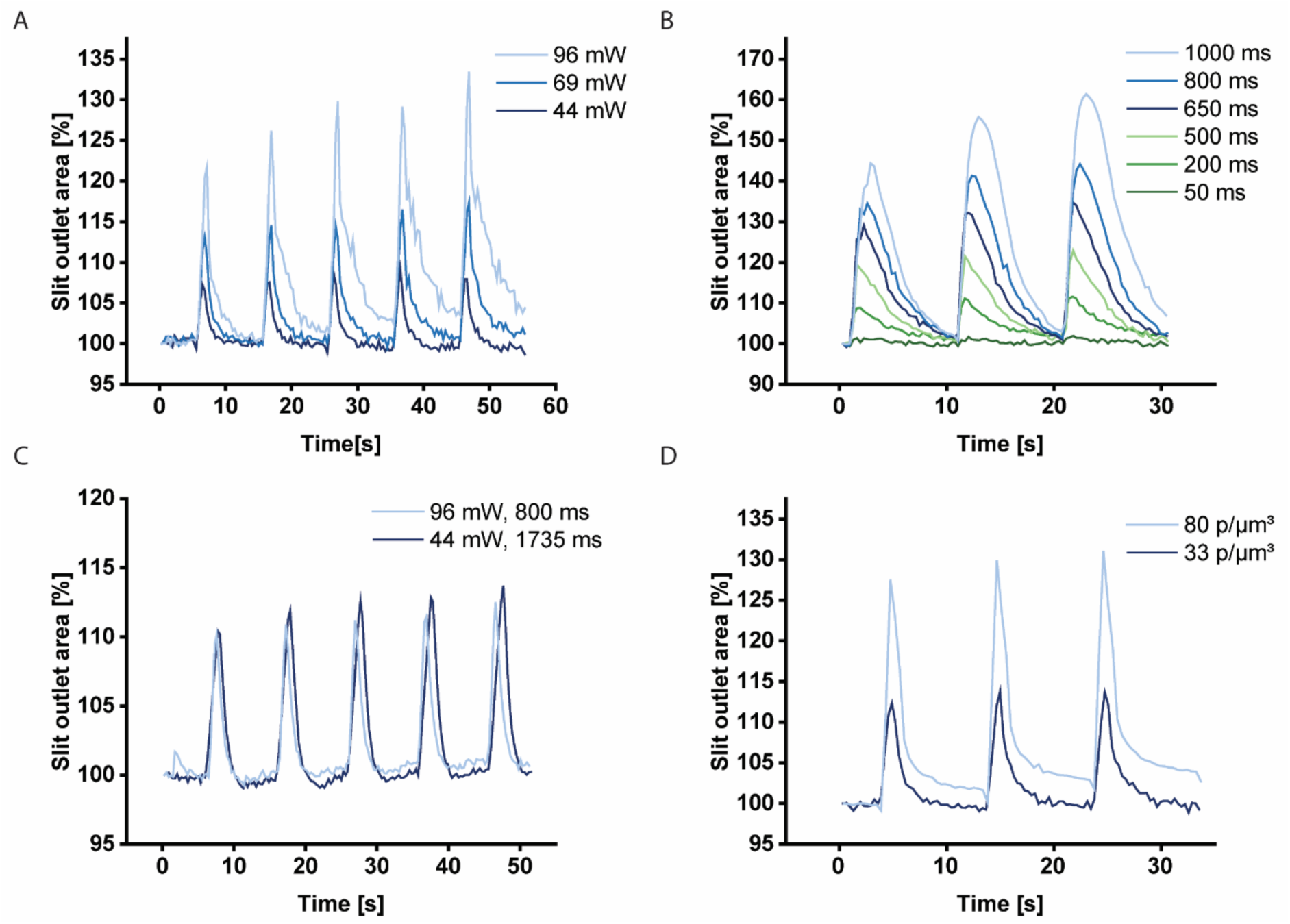
Graphs showing the effect of different parameters on the unconfined slit valve’s outlet area, specifically examining the impact of (A) laser power, (B) illumination time, (C) total laser energy per pulse, and (D) density of the AuNRs (in particles per µm^3^) in the hydrogel.

The opening of the valve can be controlled via the intensity of the laser, the duration of the exposure, and the concentration of the embedded AuNRs. Considering the results above, the properties of the laser and the gel were varied to determine how they affect the operation of the slit valve. Increasing the intensity of the laser (keeping the pulse duration constant) increases the maximum area of the valve outlet (Fig 5A). Increases in the valve outlet area between 5% and 30% are reliably obtained by varying the laser intensity from 44 mW to 96 mW. The valve opens within 1 s of the laser being turned on for all intensities used, and, for lower intensities, returns to its original size after 5-10 s. In this range, peak valve outlet areas increase approximately linearly with laser amplitude (slope = 0.3 %/mW, *R*^2^ = 0.94, as shown in Fig S6A). Note that at the highest intensities, a 10 s interval between successive illuminations is not sufficient for the valve to fully close again. This behavior can be prevented by decreasing the frequency of the laser pulses (Fig S7). Increasing the duration (on-time) of each pulse at fixed intensity similarly increases the area of the slit, up to a total of 60% for a pulse duration of 1s (Fig 5B). Peak valve outlet areas increase approximately linearly with laser amplitude (slope = 0.046 %/ms, *R*^2^ = 0.99, as shown in Fig S6B). Varying both the intensity and duration of each pulse together shows that the opening of the valve is determined by the total energy delivered by the laser. For example, 800 ms pulses at 96 mW and 1735 ms pulses at 44 mW (each delivering 76.5 ± 0.5 mJ of energy) both increase the slit area by 10-13% (Fig 5C). Lastly, increasing the density of AuNRs in the NIPAM gel from 33 particles/µm^3^ to 80 particles/µm^3^ increases the amount by which the valve opens by approximately a factor of two (Fig 5D). These results show that the operation of the photoresponsive hydrogel valve is highly tunable via several easily adjustable parameters.

**Figure 6.**
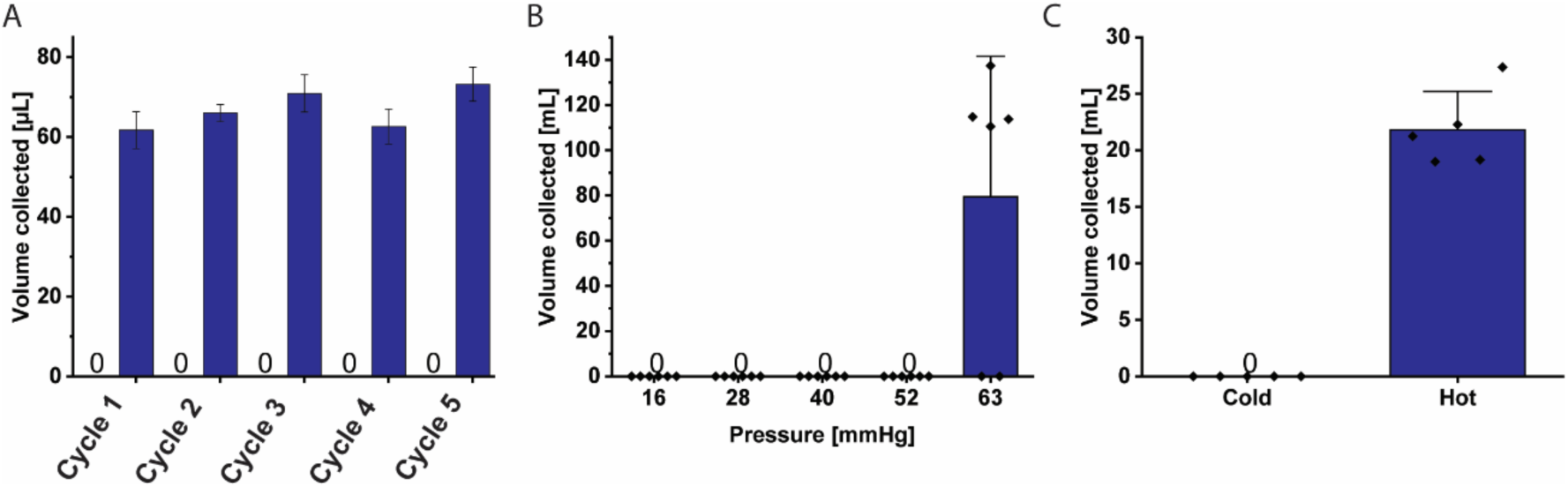
(A) Graph showing the volume of water that was collected during each cycle when the unconfined slit valve is repeatedly exposed to water at 23 °C and 80 °C. (B) Graph of the pressure resistance of the unconfined slit valve at 23 °C, showing the usable range of the valve. (C) Graph showing the flow with cold (25°C) and warm (50° C) water at glaucomatous pressure (30 mmHg). The data for the three panels were collected using 3, 6, and 5 independent replicates of the valve, respectively.

The slit valve can reversibly trigger fluid flow via a repeated change in temperature. The usability of the valve hinges upon its capacity to regulate flow consistently in response to stimuli. To further test the operation of the hydrogel valve consisting of a slit-shaped outlet in unconfined NIPAM, 200 µL of water at 23 °C and 80 °C was repeatedly added onto the valve and the volume of water passing through the valve was measured. At 23 °C, the valve does not allow any water to pass through, even after 5 cycles of exposure to cold and hot water (Fig 6A). During exposure to hot water, the slit opens considerably: between 60 µL and 80 µL of water passes through the valve during each 2-minute exposure to 80 °C water. Representative cycles at both temperatures are shown in supporting videos 2 and 3, respectively. The low variation in the volume of water collected across cycles suggests high reversibility of the valve’s thermoresponsive operation, even for very large changes in temperature (±57 °C). In practice, for biomedical applications, the valve would only have to withstand significantly smaller temperature fluctuations.

The slit valve effectively prevents the passage of water at room temperature up to at least 52 mmHg. To test the ability of the valve to withstand hydrostatic pressure at 23 °C (significantly below the VPTT), it was placed under a column of water of different heights, and the volume of water passing through the valve was measured. For pressures up to 52 mmHg (6.9 kPa), all valves tested remained impermeable, as no water was collected below (Fig 6B). At a pressure of 63 mmHg (8.4 kPa), four of the six valves allowed a large volume of water (> 100 mL) to pass, while the other two valves remained impermeable up to a maximum tested pressure of 135 mmHg (18 kPa), suggesting the design may be suitable for higher pressures with only slight adaptations.

A promising application of this valve design is the drainage of excess fluid from the eye to reduce elevated intraocular pressure (IOP), which is a major risk in glaucoma. In glaucoma patients for whom eye drops do not provide alleviation, surgically implanted devices for fluid drainage can be used. While these can lower IOP, failure rates remain significant due to post-operative complications, including both high and low IOP [9], [28], [29]. To regulate IOP with a more favorable safety profile, some new devices employ active valves to overcome the problem of hypotony (dangerously low IOP). However, a major problem is excessive scarring, bleb encapsulation, or protein adsorption and fibrosis that may lead to blocked drainage over time. The most promising solution to this problem is the use of biocompatible materials to reduce tissue reaction [9], [30], [31]. Thus, there is a need for novel active valve designs based on materials compatible with the biological surroundings to provide safe, consistent, and low-cost treatment for IOP in glaucoma patients. A miniaturized active hydrogel valve would allow excess fluid to be drained from the eye on demand while minimizing the risk of hypotony and fibrosis. Such a device would provide similar benefits to the eyeWatch^TM^ [11], but would offer greater bio- and MRI compatibility [10].

This valve design presented here can be used to control the flow of water at pressures relevant to the treatment of glaucoma. To test the thermoresponsive operation of the slit valve at a pressure of 30 mmHg (or 4 kPa – a typical intraocular pressure in patients suffering from glaucoma), valves were placed under a column of water at either 23 °C or 50 °C (well below and above the VPTT). With water at 23 °C, the valve remains completely closed and impermeable (Fig 6C). With water at 50 °C, the valve reliably opens, allowing 21.8 ± 1.5 mL of water to pass through, on average. This result demonstrates that this valve design is suitable to reliably control the flow of water at glaucomatous pressure using changes in temperature. Importantly, the hydrogel’s response to temperature can be adjusted for different physiological conditions by replacing pure NIPAM with a copolymer of NIPAM and N-ethylacrylamide (NEAM). Increasing the fraction of NEAM increases the temperature at which the gel shrinks, and thus the temperature at which the valve opens (Fig S8).

Since the slit valve operates perfectly at typical glaucomatous pressures of around 30 mmHg (and up to at least 52 mmHg), the excellent biocompatibility of NIPAM could be used to reduce complications due to fibrosis, allowing the valve to maintain its functionality in the long term. Additionally, we note that the range of hydrostatic pressures for which the valve operation was demonstrated here includes typical pressures found in many other parts of the human body, such as the bladder (10-60 mmHg) [32], [33], [34], kidney (intrarenal pressure < 20 mmHg) [35], urethra (51 mmHg and 30 mmHg maximum urethral closure pressure for continent and incontinent patients, on average) [36], and heart (4-120 mmHg on average, depending on the chamber) [37], [38].

**Figure 7.**
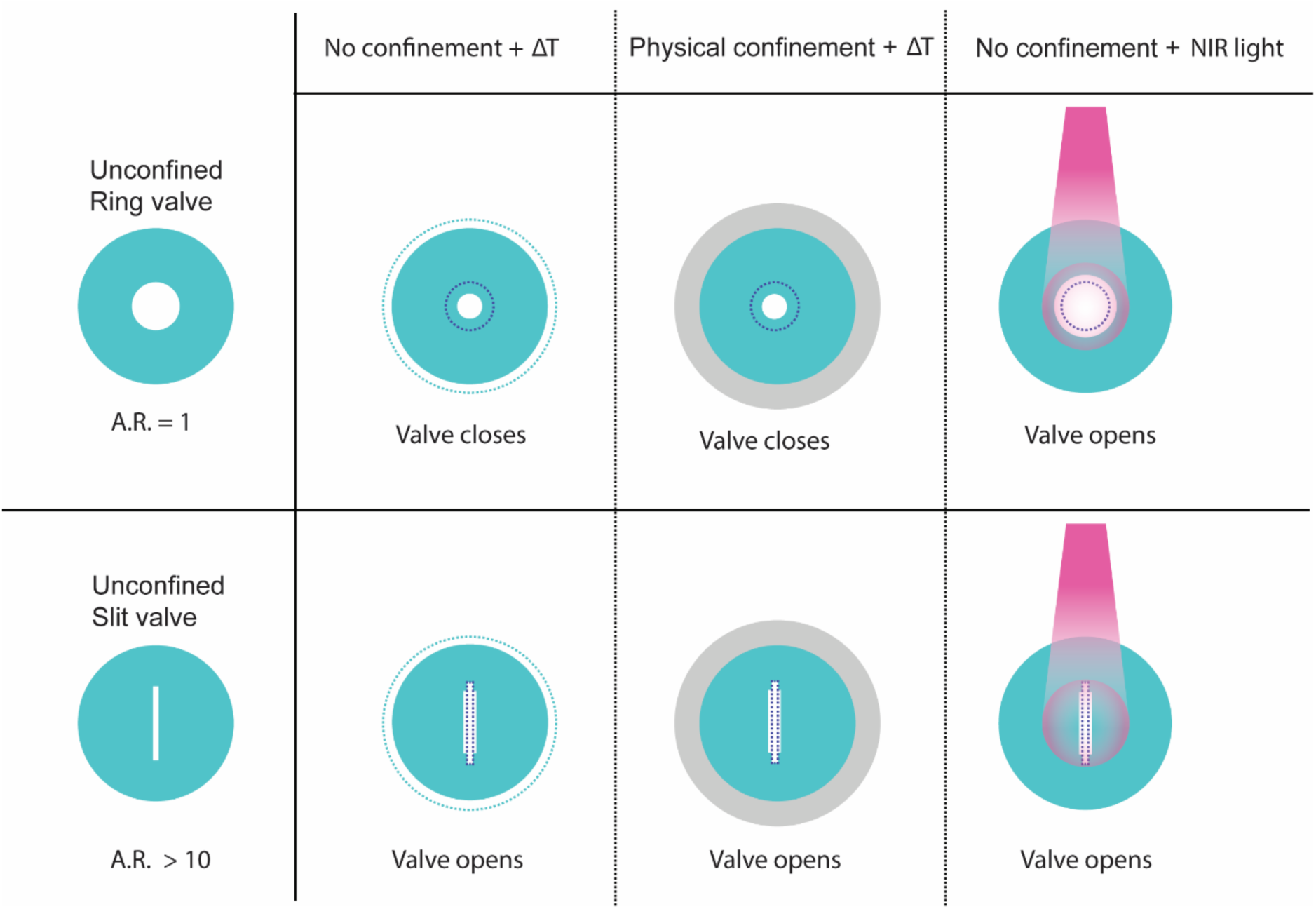
Schematic of circular and slit-shaped outlets in unconfined NIPAM valves and confined NIPAM valves, imposed by physical confinement by passive DMAA, responding to temperature and unconfined valves exposed to light. Dotted outlines represent initial dimensions.

Together, confinement of the active hydrogel and its responsiveness to both changes in temperature and exposure to light provide excellent control over the behavior of the valve (Fig 7). Circular openings in NIPAM (*AR* = 1) close in response to an increase in temperature, to a degree that can be adjusted by confining the active hydrogel with a passive one. If this same hydrogel is made photoresponsive by adding AuNRs, the valve can instead be made to open in response to localized illumination with NIR light. On the other hand, a slit-shaped opening (*R* > 10) opens both in response to an increase in temperature or to illumination with NIR light. Again, the increase in the outlet area can be adjusted via confinement, or via the parameters of the NIR illumination. Thus, this valve design provides an extremely versatile platform for the design of thermo- and photo-responsible valves for biomedical applications. It is worth noting that sufficient water must be present for the hydrogel to remain fully swelled; if it is allowed to dry out, reversible operation may be compromised.

The varied and complex changes in the size and shape of the valves can be used to show that the deformations of the hydrogel are inhomogeneous under certain conditions. In equilibrium, deformations of NIPAM hydrogels are typically believed to be homogeneous [26]. Formally, a deformation is homogeneous when the deformation gradient is independent of the spatial coordinates. Geometrically, a deformation can be shown to be homogeneous if any straight line connecting two points in the original, reference configuration of the object remains straight in the final, deformed configuration, and if any parallel lines remain parallel after the deformation. This description is consistent with the deformation observed here for a circular valve in a uniformly heated hydrogel (Fig S9). However, under any other condition described here (slit valve being uniformly heated or NIR illumination of valves of all shapes), it can be shown that certain lines do not remain straight, meaning the deformation must be inhomogeneous. Thus, the assumption that thermoresponsive hydrogels deform homogeneously does not necessarily hold when the gel is incorporated into a more complex device. Heterogeneous transformations must be considered to fully understand how such valves will behave in practice.

## CONCLUSION

The ability to control fluid flow through an active hydrogel valve depends on achieving precise control of the swelling and collapse of the outlet. Here, we show that NIPAM hydrogels can be used to fabricate active valves with highly tunable responses to both temperature and light, by varying the valve outlet’s aspect ratio, the precise method of actuation and the degree of confinement of the active gel by another, passive gel. NIPAM valves embedded with gold nanorods (AuNRs) and exposed to near infrared (NIR) light open regardless of aspect ratio, albeit to different extents. This opening behavior can be precisely modulated via the intensity, exposure time, and frequency of the incident light.

Compared to previous work with a similar design [24], this valve is about an order of magnitude smaller (1 mm), and the response of the valve is about two orders of magnitude faster, with opening times of less than 1 s and closing times of 5-20 s, depending on the laser power. These response times are over an order of magnitude faster than those of other recent designs for mm-scale active hydrogel valves [27] and are comparable to valve designs up to several orders of magnitude smaller in size [21], [28]. This improved responsiveness is due to both the precision of the NIR light stimulus used here, and the excellent photothermal properties of the AuNR-embedded NIPAM. We also show that the opening and closing of the valves is consistent across multiple cycles of illumination, as well as several independently constructed prototypes. Thus, the local confinement provided by controlled illumination of the photoresponsive NIPAM, the rapid and highly tunable behavior of the valve, and the resulting consistency of this valve’s opening and closing make it a promising candidate for a variety of applications where reliable, long-term control over fluid flow is crucial.

Future work should focus on further miniaturizing this valve design and tuning its properties to meet particular biomedical needs. A smaller valve will require even less NIR radiation intensity for its actuation and will further reduce its response time [4]. More experiments are needed to determine precisely how the pressure resistance and flow rate of the valve depend on the dimensions of the gel and the shape of the outlet at smaller scales. Additionally, the volume phase transition temperature of the active gel can be adjusted to meet the particular needs of an application. For example, the temperature at which the gel shrinks could be raised above the human body temperature by copolymerizing NIPAM with NEAM, as shown here, or by introducing a small amount (<1 %) of acrylic acid into the polymer [29], ensuring that the valve does not open except when actuated by the desired external stimulus. The precise actuation temperature should depend on the part of the body for which the device is intended (for example, the temperature on the surface of the eye tends to be slightly lower than the mean body temperature), underscoring the importance of the design presented here. In general, the particular physical properties of the active hydrogel (material strength, dimensions, elasticity, swelling ratio) can be adjusted in many ways based on the rich existing literature while taking advantage of the unique properties of local confinement, fast response times, and high tunability of the design described here. Finally, further tests of the valve’s functionality over many cycles, longer time periods, and in combination with other components or devices are warranted in the long term.

## MATERIALS AND METHODS

All materials were purchased from Sigma Aldrich, unless otherwise mentioned.

### Gel fabrication – thermoresponsive hydrogels

Unconfined NIPAM hydrogels were prepared by polymerizing a solution of 0.2 g NIPAM (99 %) in 0.341 mL dimethyl sulfoxide (DMSO, ≥99.7%) (5.2 M) with 1 mole % *N,N*′-methylenbis(acrylamide) (BIS) as crosslinker and 1 mole % lithium-phenyl-2,4,6-trimethylbenzoylphosphinate (LAP, ≥95%) as initiator. NIPAM hydrogels were fabricated by adding 43 μL of the gel precursor solution into polydimethylsiloxane (PDMS) molds with a diameter of 8 mm and a thickness of 0.85 mm. The polymerization was carried out under UV light (365 nm) at 2.5 cm distance from the hydrogel surface, with nitrogen (N_2_) purging for 5 min to minimize oxygen contact during polymerization process. Fluorescent NIPAM gels were prepared by adding 0.4 mg of methacryloxyethyl thiocarbamoyl rhodamine B (Polysciences Inc) to 100 μL of NIPAM precursor solution.

Confined hydrogels were fabricated using the same NIPAM precursor solution, with an additional surrounding *N,N*-dimethylacrylamide (DMAA) solution prepared from 2 M DMAA in DMSO, containing 0.04 M BIS as a crosslinker and 4 mg/mL LAP as a photoinitiator. The protocol followed that of previous work [24], which showed that partial polymerization of one hydrogel (DMMA) followed by the addition and subsequent polymerization of a second gel (NIPAM), results in a strongly bonded interface between the two gels. Here, the same cross-linker (BIS) was used for both gels, ensuring cross-linking at the interface during the second polymerization step. Ring-shaped PDMS molds of different sizes (14 and 20 mm diameter) and a PDMS disc (8 mm diameter) in the center as a placeholder for the NIPAM part of the confined hydrogel were attached to glass slides (Marienfeld) using plasma (high vacuum) for 120 seconds. 88 µL (14mm) or 224 µL (20mm) of DMAA solution were pipetted in this mold, and partially polymerized for 5 min. The center PDMS disc was then manually removed and 43 µL of NIPAM precursor solution were added to the cavity and polymerized for 5 min with UV with N_2_ purging. After polymerization the outer PDMS mold was removed from the glass slide before the hydrogels were swelled overnight in excess water to remove all DMSO.

### Preparation of valve outlets

Valve prototypes were prepared by creating an outlet in the swelled hydrogel using a biopsy punch (O.D. = 1 mm) for circular-shaped outlet and beaver blades (0.1 mm thickness) that were laser-cut (Lasercut AG) to 1 mm length for the slit-shaped outlet. The valves were always produced on polystyrene substrate, as this material provided the sharpest borders of the valve. The punch was used manually, whereas the blade was fitted on a customized press (Fig S10). Due to high variability of the slit and hole production, specific inclusion criteria were established. For slits, the criterion was a clear distinction of the slit area from the surrounding material, while for holes, it was an intact hole without cracks at the outlet. For this, all the outlets were checked visually under the microscope to ensure functionality and were excluded if they did not meet the inclusion criteria.

### Thermal measurement settings

NIPAM is a thermoresponsive hydrogel, collapsing at a specific volume phase transition temperature (VPTT), usually 34 °C. Therefore, we were interested in the outlet behavior as well as the NIPAM diameter at temperatures between 25 °C and 40 °C. For this, the hydrogels were placed in a petri dish (diameter = 30 mm) containing 3 mL water, which was placed on a customized heating mat (50 W, silicone, RS Pro). The heating mat was connected to a temperature sensor (PT100, platinum, Innovative Sensor Technology (iST)) via a temperature controller (ET2011, Enda) to monitor and set the temperature accurately. The sensor was positioned at the edge of the petri dish, fully submerged in the water without contact with the dish. Additionally, the experiment was conducted in a heating chamber (customized) for the best possible temperature stabilization. The heating chamber and the equilibrium time of 5 min ensured that there was no temperature gradient of the water where the hydrogels were placed as well as no difference in temperature between confined and unconfined hydrogels. The water temperature in the petri dishes for both confined and unconfined samples was measured before each experiment, verifying that it remains consistent between unconfined and confined hydrogels with 5 min equilibrium time in the heating chamber.

### Thermal response – bulk hydrogels

The diameter of NIPAM hydrogels in both confined and unconfined conditions was measured at various temperatures to evaluate the influence of physical confinement on hydrogel collapse. Images were acquired using a NIKON camera equipped with a 105 mm objective. All images were analyzed and quantified using Fiji. The hydrogel diameter was manually measured in Fiji, and the values were normalized to the diameter of the respective confinement group at 25 °C to account for sample variability.

### Thermal response – copolymer hydrogels

The diameter of unconfined hydrogels consisting of different copolymers of NIPAM and NEAM (99%) was measured to evaluate the effect of chemical composition on the VPTT. Hydrogels with an initial diameter of 8 mm were polymerized and allowed to swell overnight in MQ water. Subsequently, 3 mm diameter hydrogels were punched out using a biopsy punch. Each sample was immersed in 200 µL of MQ water and incubated at different temperatures (25, 28, 30, 32, 34, 37, 40, and 50 °C) using a customized heating mat. At each temperature, the hydrogel diameter was measured and normalized to its diameter at 25 °C to assess relative swelling behavior.

### Thermal response – valve outlets

The outlet area of unconfined and confined (20 mm) hydrogels with either a 1 mm slit or 0.5 mm diameter circular outlet was measured at varying temperatures. For image acquisition, an Axio Imager M1 inverted fluorescence microscope (Zeiss) with a 4x Objective was used. Images were captured using both brightfield and fluorescence with excitation and emission wavelengths of 543 nm and 565 nm, respectively, for the rhodamine. All images were analyzed and quantified using Fiji (ImageJ Java 8), with the image scale set to 0.6202 pixels/µm.

For the analysis of the circular outlet, a macro was applied to the rhodamine channel image using Huang Auto thresholding. Outlet areas were identified and measured using the “Analyze Particles” tool, with circularity of 0.5 to 1 and a size of 60,000–2,000,000 µm². The analysis provided the thresholded image, an outline drawing of the detected outlet, and the calculated outlet area (Fig S11). In cases where the macro did not perform adequately, the outlet area was manually measured using polygon selections.

For the slit outlets, measurements were performed manually on the brightfield channel using polygon selections to outline and measure the outer area of the slit (Fig S12). Due to the morphological differences between the two outlet types—where the circular outlet remains open at all times, while the slit outlet opens only at higher temperatures—it was not possible to analyze both using the same macro approach. In all cases, the measured outlet areas were normalized to the outlet area at 25 °C to account for variations across samples and conditions.

### Reversibility of valve action

The reversibility of the opening and closing of the slit-shaped outlet was tested in an unconfined NIPAM hydrogel (n=3) over five cycles. A cycle was defined as one measurement at low temperature (23 °C water) followed by one measurement at high temperature (80 °C water). The swollen hydrogel with a 1 mm slit was placed on a petri dish with a 4 mm diameter hole, and a beaker was positioned underneath to collect any liquid flowing through the valve. The slit location on the petri dish hole was checked visually before and after each measurement.

First, 200 µL of methylene blue (MB) stained water (0.005 g/L) at 23 °C were pipetted onto the hydrogel. After 2 minutes, the weight of the collecting beaker was measured before the hydrogel was allowed to swell in 23 °C water for 2 minutes. Subsequently, the hydrogel was placed back on the petri dish, and 200 µL of 80 °C MB-stained water was pipetted onto the hydrogel. After waiting for 2 minutes to allow the valve to open, the weight of the collecting beaker was measured again. The hydrogel was then allowed to swell in water for another 2 minutes to close the hydrogel, following the same procedure as for the room temperature water.

This process was repeated for all hydrogels five times (cold–warm cycles). The pipetting process, as well as the 2-minute waiting times, were filmed to provide proof of the procedure.

### Resistance to hydrostatic pressure

The ability of the hydrogel valve to withstand hydrostatic pressure is relevant for applications such as glaucoma or urinary valves. Confined slit-shaped valves (n=6) were subjected to relevant pressure using a custom-made setup (Fig S13) where an aluminum column (I.D. = 20 mm, l = 2 m) was placed on the DMAA part of the hydrogel. To prevent destruction of the hydrogel, a PDMS ring (T= 0.85 mm, O.D. = 30 mm, I.D. = 18 mm) was placed between hydrogel and column. The slit valve was aligned with the hole in the petri dish and the alignment was checked before and after the pressure measurement. The column was then filled with 23°C water using a peristaltic pump (Watson Marlow 400, 403 U/RI, 50 RPM) at maximum speed, waiting 1 minute at each pressure before increasing it. The collecting beaker was weighed at each pressure. Each hydrogel was measured until the first leakage was observed. The hydrostatic pressure was calculated using the following equation

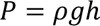

Where *P* = hydrostatic pressure in mmHg, *ρ* = density of water, *g* = gravitational constant and *h* = height of water in the aluminum column.

Additional control experiments were performed with hydrogels without a valve, which completely prevented flow into the beaker for all pressures tested, confirming that any liquid collected had passed through the valve outlet.

### Flow measurement

The same customized setup used for testing resistance to hydrostatic pressure was employed, using a different aluminum column (I.D. = 20 mm, l = 45 cm). The confined slit-shaped valve (n=5) was subjected to a glaucomatous pressure of 30 mmHg (4 kPa) to demonstrate its stability at this pressure. The column was filled with 23°C water using a peristaltic pump (Watson Marlow 400, 403 U/RI, 50 RPM) at maximum speed. After waiting for 5 minutes, the weight of the collecting beaker was measured and the water was carefully removed from the column using the peristaltic pump. To measure the water flux when the valve was open, warm water (50°C) was used. It was ensured that the water temperature in the column remained above 34°C throughout the 5 min measurement. After 5 minutes, the flow volume was measured on the balance.

### Preparation of AuNRs

The AuNRs used in this study were synthesized by a seed-mediated growth method from chloroauric acid (HAuCl_4_) and functionalized with thiol-poly(ethylene glycol) (PEG-SH) to stabilize and prevent aggregation, as previously described elsewhere [30]. 50 mL HAuCl_4_ solution (0.003 M) were mixed with 100 mL Milli-Q water and stirred at 25°C (250 – 300 rpm). 150 mL of cetyltrimethylammonium bromide (CTAB, 0.2 M) solution (VWR International) were added very slowly with stirring to avoid foaming. After stirring for 20 – 30 min, 3.1 mL ascorbic acid (Fluka) solution (0.05 M) were added dropwise. Then 3.3 mL of AgNO_3_ solution (0.008 M) were added and stirred continuously to promote the formation of Au(I) (growth solution).

In a second flask, 4.2 mL of iced Milli-Q water, 5 mL of CTAB solution, and 830 µL of HAuCl_4_ solution were mixed with stirring. 600 µL of freshly prepared, cold NaBH_4_ solution (0.01 M) were quickly added and stirred at 1200 rpm for 2 min. Then the solution was stirred for another 2 min at 700 rpm and left for 30 min without stirring to achieve seed formation (seed solution). 875 µL of seed solution were added to the growth solution while stirring at 1200 rpm. After 5 min, the prepared solution was stirred at 250 – 300 rpm for 30 min. Then, 2 mL of ascorbic acid solution were added once the solution turned purple using an Aladdin syringe pump (AL-1000, World Precision Instruments) (500 µL/h) with gentle stirring for 4 h to reduce Au(III) and form AuNRs. Stirring was continued for further 30 min.

After 4.5 h, the solution was centrifuged (35 mL/centrifuge tube) at 9500 rpm at room temperature (RT) for 20 min and the supernatant was removed. The concentrated AuNRs were filtered with a polytetrafluoroethylene (PTFE) filter (0.2 µm) and diluted with 20 mL milli-Q water.

The AuNRs were modified with PEG after synthesis to prevent aggregation. For this purpose, 32 mg of PEG-SH (Iris Biotech) and a few Tris (2-carboxyethyl) phosphine (TCEP) crystals (to prevent the formation of disulfide bridges) were added to a solution of AuNRs in 20 % (v/v) ethanol in water. The solution was then ultrasonicated first at 60°C for 30 min and then at 20°C for 3.5 h, after which the AuNRs were transferred to DMSO, concentrated and added to pre-polymer solutions. The optical density of the AuNR in DMSO was measured with a nanodrop instrument (SynergyH1_Biotek) and the pegylation was proofed by TEM (Fig S14).

### Transmission electron microscopy

AuNR were evaluated using a Zeiss EM900 Transmission electron microscopy (TEM). Holey carbon supported copper grid (300 Mesh, 5 nm diameter) was used as a substrate. The concentrated gold nanorod solution was diluted 1: 25’000 in DMSO. A volume of 5 µL of the diluted gold nanorod solution was applied to the grids and incubated for 1 min. After incubation, the grids were dried before TEM analysis was performed. The aspect ratio, important for the absorbance of the AuNR, as well as the pegylation was evaluated (Fig S14).

### Gel fabrication – light responsive hydrogels

PEG-functionalized AuNR in DMSO were incorporated into the NIPAM prepolymer solution with a density of 33 particles/µm^3^ (unless stated otherwise). The hydrogel production is comparable to the one without AuNR. The difference is in the UV polymerization of the NIPAM prepolymer with AuNR where an increased distance between hydrogel and UV source of 9 cm is used. This is done to prevent the heating of the AuNR during the polymerization, which leads to AuNR aggregation and therefore pattern formation in the hydrogel.

### NIR irradiation

An Axio Imager M1 (Zeiss) inverted fluorescence microscope was customized in-house to render it suitable for irradiation of samples with NIR light (Fig S4A). For this purpose, a diode laser (Roithner Lasertechnik RLTMDL-800-500-3, λmax 800 nm, max power 2’500 mW) was connected to a fiber optic cable (400 µm) and a focus-adjustable collimator (SMA905). The output power at 100% was adjusted to 480 mW (Class 3B) by replacing the resistance at R56/R96 with 500 Ω using a potentiometer. The collimator was held in place using an arm that allowed for alignment of the laser. The laser beam was aligned so that the beam was visible through the microscope and the valve (slit and circular outlet) was centered and completely in the beam of the laser (Fig S4B). The laser footprint was determined before every experiment and was approximately 1.5 mm^2^ (1.4 mm diameter). This laser setup enabled continuous laser irradiation with adjustable laser amplitudes. Using an additional customized external controller, the laser could be operated not only in continuous mode but also in pulsed mode, allowing variation of both the laser interval and on time. The output of the laser (mW) in response to the applied forward bias current was determined using a power meter (Coherent PM2) with a detector (Coherent FieldMax II) (Fig S15) and the area of the footprint was used to calculate the intensity of the laser. In this study, laser intensities ranging from 29 to 102 kW/m^2^ were used. The short pass filter (Edmund Optics OD4 775 nm) was used to protect the AxioCam from the laser beam. The opening and closing of the valve outlets were captured using an Axio Imager M1, with time series acquired at a frame rate of approximately 0.3 frames per second using AxioCam MRm in streaming mode.

### Image analysis of light responsive outlets

Valve outlet sizes were quantified from brightfield channel images using Fiji (ImageJ Java 8). Movies showing the morphological changes to the valve outlet shape and size in response to NIR irradiation were analyzed using a macro (Fig S13), which performed the following steps: background subtraction, Otsu thresholding minimizing the intra-class variance, and particle analysis to filter objects by size (5’000-500’000 μm^2^) and circularity (0.5-1 for circular outlet and 0-0.3 for slits). The area of the valve outlet was used to define valve kinetics in response to NIR light. Three replicates (Fig S6) were tested, one of which was plotted in Fig 5.

### Kinetics – laser parameters and analysis

Characterizing the kinetics of the valve outlet is essential to precisely evaluate the performance of the light-activated valve. Since these kinetics are directly influenced by laser parameters such as amplitude, frequency and on-time as well as the density of AuNRs within the hydrogel, we investigated the impact of these parameters in an unconfined hydrogel with a 1 mm slit outlet. The hydrogel was placed on a glass substrate surrounded by a PDMS ring serving as a mold, which was filled with 1 mL of Milli-Q water to prevent drying of the hydrogel.

All experiments involving laser parameter variations were conducted on the same hydrogel sample (except for the AuNR density tests) to minimize variations in slit morphology. The influence of laser amplitude was assessed by keeping the interval time constant at 10’000 ms (0.1 Hz) and the on time at 800 ms, while testing laser powers of 44, 69, and 96 mW. Additional experiments were performed at a fixed laser energy of 77 mJ. The effect of laser pulse duration on valve operation was studied by varying the on-time between 50 and 1’000 ms, while keeping the interval time at 10’000 ms (0.1 Hz) and the laser power at 96 mW. For the frequency experiments, the laser power was set to 96 mW with an on time of 500 ms, and the interval time was varied from 2’000 to 20’000 ms (frequencies between 0.5 and 0.05 Hz). To assess the influence of AuNR density, a hydrogel with 33 particles/µm^3^ was compared to a hydrogel with 80 particles/µm^3^.

**Table 1.**
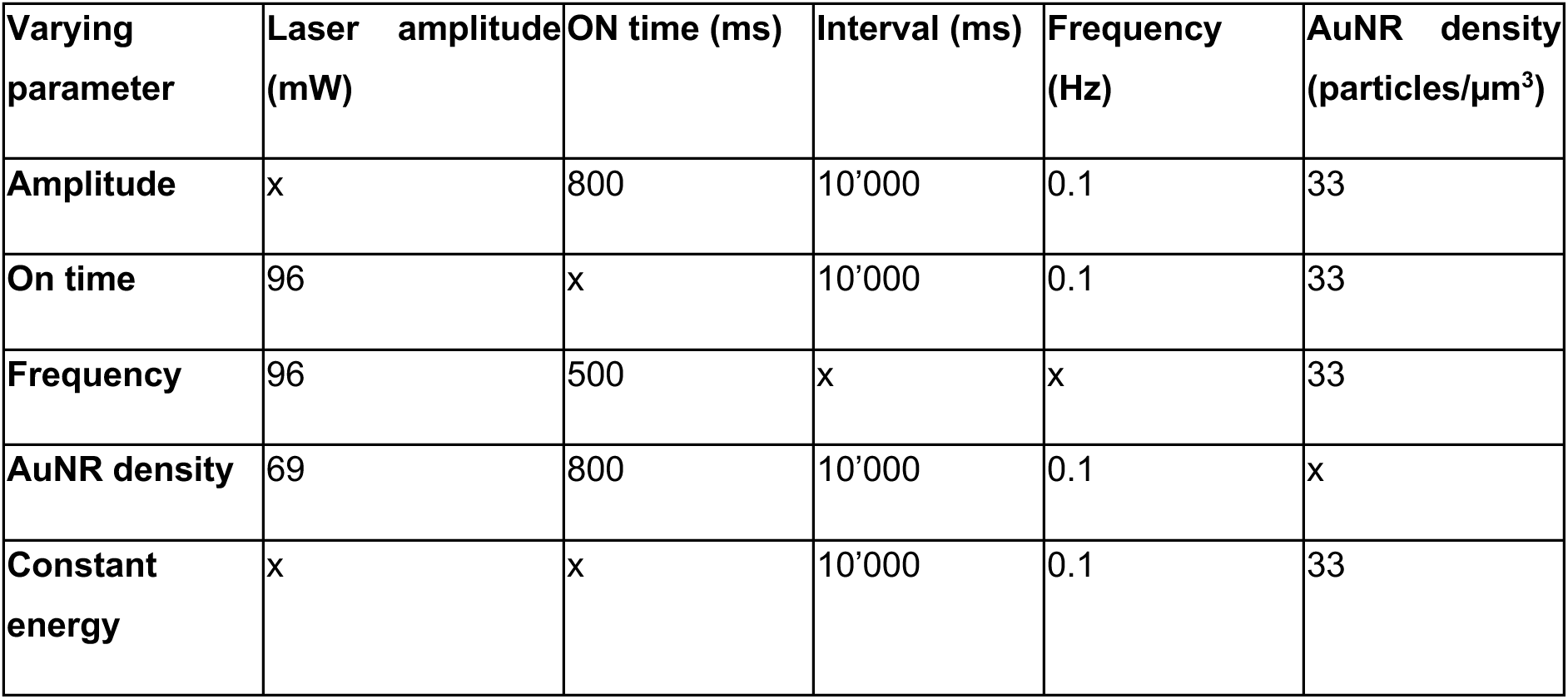
Overview of the experimental conditions used to investigate the influence of the laser parameters and AuNR density in the hydrogel on the valve outlet kinetics. For each evaluation, one parameter was varied (indicated by x) while the others were kept constant.

Image analysis was performed using a macro with the following processing steps. First, the image was cropped and the background subtraction (light background) was applied using a rolling ball radius of 50 pixels. Then, automatic thresholding was performed using the Otsu method. The slit outlet was analyzed using the “Analyze Particles” function, with a size threshold of 30’000 µm² to infinity and a particle circularity range of 0 to 0.5. For each identified particle, the outlines were displayed, and the area within the outline was measured. The slit outlet area was then normalized to the original slit outlet area of each time series.

### Statistical analysis

Statistical analysis was done in GraphPad Prism (version 10.4.2). Values are presented as mean ± standard error, unless otherwise mentioned. Each experiment is repeated at least 3 times in the form of 3 independent experiments containing a minimum of 3 samples per condition per time point, unless otherwise stated. The performed tests with relevant significances are described in the figure legends. Statistical significance was determined using a one-way or two-way ANOVA followed by Tukey multiple comparison tests. All reported p-values are multiplicity-adjusted to take into account the multiple comparisons performed, and *, **, *** correspond to p < 0.05, 0.01 and 0.001, respectively.

### Data availability

All original measurements, raw images, analysis, and imageJ macros are made available on Zenodo at https://doi.org/10.5281/zenodo.17272468.

## ACKNOWLEDGEMENTS

Technical support from Jörg Gschwend (Empa), Stefan Gfeller (Empa) and Peter Huber (Empa) is greatly appreciated. Y.C and A.M are grateful to Dr. Martina Viola (Empa) for support with image illustration for Figure S13 and to Dr. Vera Kissling (Empa) for support with TEM image acquisition. Financial support from the Heinz A. Oertli Fonds, the LHW Stiftung, the Gemeinnützige Stiftung für medizinische hilfe and the Hedy-Glor-Meyr Stiftung and the Empa Zukunftsfonds is acknowledged.

## AUTHOR CONTRIBUTIONS

A.M.: Conceptualization, Methodology, Formal Analysis, Investigation, Visualization, Writing – Original Draft, Writing – Review & Editing.

V.H.: Methodology, Formal Analysis, Investigation, Visualization, Writing – Original Draft, Writing – Review & Editing.

K.M: Conceptualization, Writing – Original Draft, Writing – Review & Editing, Funding acquisition. L.B.: Conceptualization, Writing – Review & Editing, Funding acquisition.

R.R.: Conceptualization, Writing – Review & Editing.

M.R.: Conceptualization, Methodology, Supervision, Writing – Original Draft, Writing – Review & Editing, Funding acquisition.

Y.C.: Conceptualization, Methodology, Formal Analysis, Investigation, Visualization, Supervision, Writing – Original Draft, Writing – Review & Editing, Funding acquisition.

