## Supporting Information for "Towards light responsive hydrogel-based valves for flow regulation"

**Supporting video 1:**

*Real-time video of confined slit valve opening under NIR illumination. For this recording, laser power was set to 95%, on-time 2s, and frequency 0.17 Hz.*

**Supporting videos 2 and 3:**

*Real-time videos of an experiment described in Fig 6, showing unconfined slit valves exposed to water at 23 and 80 °C, respectively. The videos are consecutive and show the same valve.*

**Supporting figures:**

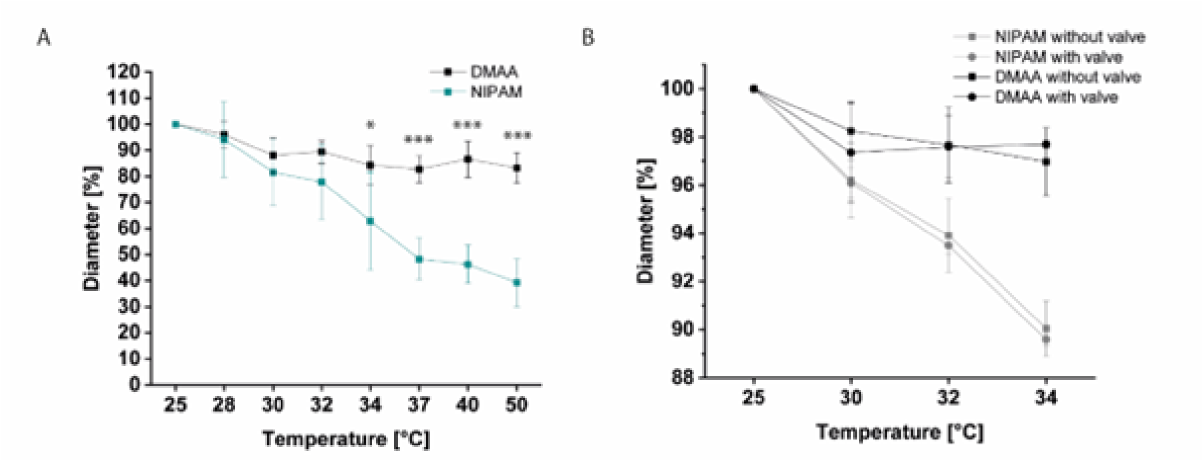

Figure S1. Experimental setup for determining the transition temperatures of DMAA and NIPAM hydrogels. Hydrogels (8 mm diameter) were prepared, swollen overnight, and smaller samples (n = 6) were obtained using a 3 mm biopsy punch. The samples were placed in a 48-well plate with 200 µL Milli-Q water per well. Temperature control was achieved using a customized heating mat (25–50 °C). At each temperature step, hydrogel dimensions were recorded using a NIKON camera with a 105 mm objective. Image analysis and manual diameter measurements were performed in Fiji to assess temperature-dependent swelling behavior. Thermal response of A) DMAA and NIPAM hydrogels showing no notable thermal response of DMAA but a significant diameter reduction of NIPAM with a volume phase transition temperature (VPTT) between 32 and 34 °C.* indicates the significance withing the material group B) Investigation of the influence of the outlet on the diameter of NIPAM and DMAA hydrogels. Confined hydrogels with a 1mm hole outlet (n = 3) and without an outlet (n = 3) were placed in a 30 mm Petri dish containing 3 mL of Milli-Q water. Diameters were measured at 25, 30, 32, and 34 °C. For imaging, hydrogels were briefly transferred into a 60 mm Petri dish without Milli-Q water to improve visualization, particularly of DMAA samples, before being returned to the original dish to prevent dehydration. Manual diameter measurements were performed using Fiji. Statistical analysis revealed no significant differences between samples with and without an outlet, indicating that the presence of the outlet does not affect the diameter of NIPAM or DMAA hydrogels. confined hydrogels (NIPAM and DMAA measured) without and with valve. *, **, *** were determined using one-way ANOVA and are statistically significant at $p < 0.05$, 0.01 and 0.001, respectively.

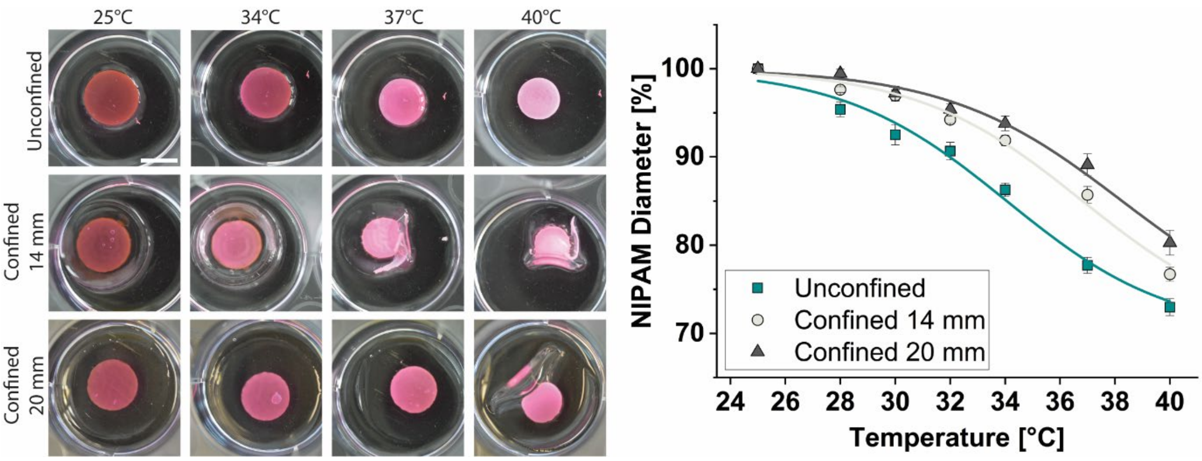

*Figure S2. Left: Photos showing the unconfined, and confined gels (without outlet), with 2 different degrees of confinement (14 mm and 20 mm). Scale bar 1 cm. Right: Diameter of unconfined and confined hydrogels as a function of temperature (reproduced from Fig 2). Solid lines represent logistic fits according to the equation* $D = A - \frac{B}{1+e^{-k(T-T_0)}}$ *with fitting parameters shown in Table S1.*

| | **A** | **B** | **k** | **$T_0$** | **Adjusted $R^2$** |
|---|---|---|---|---|---|
| **Unconfined** | 100 | 30 | 0.334 | 34.06 | 0.98 |
| **Confined 14 mm** | 100 | 30 | 0.330 | 36.82 | 0.99 |
| **Confined 20 mm** | 100 | 30 | 0.314 | 38.26 | 0.98 |

Table S1. Fitting parameters for logistic fits to NIPAM diameters as functions of temperature (see Fig S2). Parameters A and B were chosen manually and fixed to allow the fits to converge and to avoid overfitting. The remaining parameters k and *$T_0$ describe the overall rate of change of the diameter near the transition and the temperature at which the rate of change appears to be highest, respectively.*

| **Two-way ANOVA** | | | | | |
|---|---|---|---|---|---|
| Alpha | 0.05 | | | | |
| **Source of Variation** | **% of total variation** | **P value** | **P value summary** | **Significant?** | |
| Interaction | 2.906 | <0.0001 | **** | Yes | |
| Temperature | 84.18 | <0.0001 | **** | Yes | |
| Confinement | 8.248 | <0.0001 | **** | Yes | |
| **ANOVA table** | **SS** | **DF** | **MS** | **F (DFn, DFd)** | **P value** |
| Interaction | 250 | 12 | 20.84 | F (12, 105) = 5.454 | P<0.0001 |
| Temperature | 7244 | 6 | 1207 | F (6, 105) = 316.0 | P<0.0001 |
| Confinement | 709.7 | 2 | 354.9 | F (2, 105) = 92.89 | P<0.0001 |
| Residual | 401.1 | 105 | 3.82 | | |
| **Data summary** | | | | | |
| Columns (Confinement) | 3 | | | | |
| Rows (Temperature) | 7 | | | | |
| Number of values | 126 | | | | |

*Table S2. Two-way ANOVA results for hydrogel disk diameters as functions of both temperature and confinement (data shown in Fig 2 and Fig S2). Analysis performed using GraphPad Prism.*

| Tukey's multiple comparisons test | Mean diff. | 95.00% CI of diff. | Summary | Adjusted P Value |
|---|---|---|---|---|
| 25 | | | | |
| Unconfined vs. Confined 14 mm | 0 | -2.173 to 2.173 | ns | >0.9999 |
| Unconfined vs. Confined 20 mm | 0 | -2.173 to 2.173 | ns | >0.9999 |
| Confined 14 mm vs. Confined 20 mm | 0 | -2.173 to 2.173 | ns | >0.9999 |
| **28** | | | | |
| Unconfined vs. Confined 14 mm | -2.235 | -4.408 to -0.06216 | * | 0.0423 |
| Unconfined vs. Confined 20 mm | -4.07 | -6.243 to -1.897 | **** | <0.0001 |
| Confined 14 mm vs. Confined 20 mm | -1.835 | -4.008 to 0.3381 | ns | 0.1154 |
| **30** | | | | |
| Unconfined vs. Confined 14 mm | -4.43 | -6.603 to -2.257 | **** | <0.0001 |
| Unconfined vs. Confined 20 mm | -4.687 | -6.860 to -2.514 | **** | <0.0001 |
| Confined 14 mm vs. Confined 20 mm | -0.257 | -2.430 to 1.916 | ns | 0.9574 |
| **32** | | | | |
| Unconfined vs. Confined 14 mm | -3.565 | -5.738 to -1.392 | *** | 0.0005 |
| Unconfined vs. Confined 20 mm | -4.746 | -6.919 to -2.572 | **** | <0.0001 |
| Confined 14 mm vs. Confined 20 mm | -1.18 | -3.353 to 0.9930 | ns | 0.4033 |
| **34** | | | | |
| Unconfined vs. Confined 14 mm | -5.602 | -7.775 to -3.429 | **** | <0.0001 |
| Unconfined vs. Confined 20 mm | -7.514 | -9.687 to -5.341 | **** | <0.0001 |
| Confined 14 mm vs. Confined 20 mm | -1.912 | -4.085 to 0.2612 | ns | 0.0965 |
| **37** | | | | |
| Unconfined vs. Confined 14 mm | -7.987 | -10.16 to -5.814 | **** | <0.0001 |
| Unconfined vs. Confined 20 mm | -11.39 | -13.56 to -9.216 | **** | <0.0001 |
| Confined 14 mm vs. Confined 20 mm | -3.402 | -5.576 to -1.229 | *** | 0.0009 |
| **40** | | | | |
| Unconfined vs. Confined 14 mm | -3.732 | -5.905 to -1.559 | *** | 0.0003 |
| Unconfined vs. Confined 20 mm | -7.305 | -9.478 to -5.132 | **** | <0.0001 |
| Confined 14 mm vs. Confined 20 mm | -3.573 | -5.746 to -1.400 | *** | 0.0005 |

*Table S3. Results of post-hoc test following the ANOVA in Table S2. Tukey multiple comparisons tests were performed to test whether the degree of confinement was significant at each temperature. Multiplicity-adjusted p-values are reported to account for multiple comparisons.*

| Two-way ANOVA | Ordinary |
|---|---|
| Alpha | 0.05 |

| Source of Variation | % of total variation | P value | P value summary | Significant? |
|---|---|---|---|---|
| **Circular valve** | | | | |
| Interaction | 11.14 | <0.0001 | **** | Yes |
| Temperature | 53.71 | <0.0001 | **** | Yes |
| Confinement | 13.35 | <0.0001 | **** | Yes |
| **Slit valve** | | | | |
| Interaction | 6.727 | 0.0183 | * | Yes |
| Temperature | 49.69 | <0.0001 | **** | Yes |
| Confinement | 3.580 | 0.0068 | ** | Yes |

| ANOVA table | SS (Type III) | DF | MS | F (DFn, DFd) | P value |
|---|---|---|---|---|---|
| **Circular valve** | | | | | |
| Interaction | 2260 | 5 | 451.9 | F (5, 86) = 25.53 | P<0.0001 |
| Temperature | 10899 | 5 | 2180 | F (5, 86) = 123.2 | P<0.0001 |
| Confinement | 2709 | 1 | 2709 | F (1, 86) = 153.1 | P<0.0001 |
| Residual | 1522 | 86 | 17.70 | | |
| **Slit valve** | | | | | |
| Interaction | 593690 | 5 | 118738 | F (5, 90) = 2.887 | P=0.0183 |
| Temperature | 4385755 | 5 | 877151 | F (5, 90) = 21.33 | P<0.0001 |
| Confinement | 315968 | 1 | 315968 | F (1, 90) = 7.683 | P=0.0068 |
| Residual | 3701227 | 90 | 41125 | | |

| Difference between column means | Circular valve | Slit valve |
|---|---|---|
| Predicted (LS) mean of Unconfined | 81.29 | 219.1 |
| Predicted (LS) mean of Confined | 91.95 | 330.6 |
| Difference between predicted means | -10.66 | -111.5 |
| SE of difference | 0.8615 | 40.23 |
| 95% CI of difference | -12.37 to -8.946 | -191.4 to -31.59 |

| Data summary | Circular valve | Slit valve |
|---|---|---|
| Number of columns (Confinement) | 2 | 2 |
| Number of rows (Temperature) | 6 | 6 |
| Number of values | 98 | 102 |

*Table S4. Two-way ANOVA results for valve outlet areas as functions of both temperature and confinement (data shown in Fig 3). Values for circular and slit valve are shaded blue and green, respectively. Analysis performed using GraphPad Prism.*

| Tukey's multiple comparisons test | Predicted (LS) Mean diff. | 95.00% CI of diff. | Summary | Adjusted P Value |
|---|---|---|---|---|
| Unconfined vs. Confined | | | | |
| **Circular valve** | | | | |
| 25 | 0.000 | -4.064 to 4.064 | ns | >0.9999 |
| 28 | -0.8489 | -4.913 to 3.215 | ns | 0.6790 |
| 30 | -5.332 | -9.396 to -1.268 | * | 0.0107 |
| 32 | -11.07 | -15.13 to -7.007 | **** | <0.0001 |
| 34 | -17.84 | -22.06 to -13.63 | **** | <0.0001 |
| 37 | -28.86 | -33.52 to -24.19 | **** | <0.0001 |
| **Slit valve** | | | | |
| 25 | 0.000 | -195.8 to 195.8 | ns | >0.9999 |
| 28 | 3.200 | -192.6 to 199.0 | ns | 0.9742 |
| 30 | -8.993 | -204.8 to 186.8 | ns | 0.9275 |
| 32 | -49.77 | -245.5 to 146.0 | ns | 0.6147 |
| 34 | -197.1 | -392.9 to -1.355 | * | 0.0485 |
| 37 | -416.4 | -612.1 to -220.6 | **** | <0.0001 |

*Table S5. Results of post-hoc test following the ANOVA in Table S4. Tukey multiple comparisons tests were performed to test whether the degree of confinement was significant at each temperature. Multiplicity-adjusted p-values are reported to account for multiple comparisons. Values for circular and slit valve are shaded blue and green, respectively.*

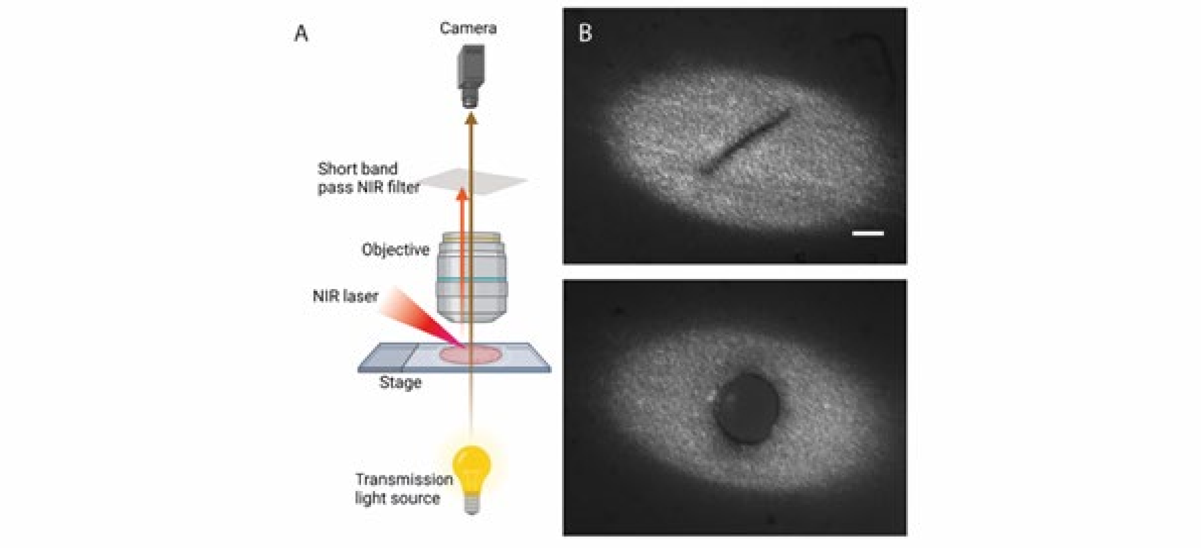

*Figure S3. (A) The hydrogel sample is irradiated with near-infrared (NIR) light (shown in orange) directed through the microscope objective. A short-pass NIR filter is placed in the detection path to protect the camera by blocking NIR wavelengths while allowing visible light (brown) to pass through. (B) The images show the centralized alignment of the laser spot on the outlet. The laser spot size (~1.5 mm²) is sufficiently large to cover the entire outlet area, ensuring uniform irradiation across the hole. Scale bar 200 µm*

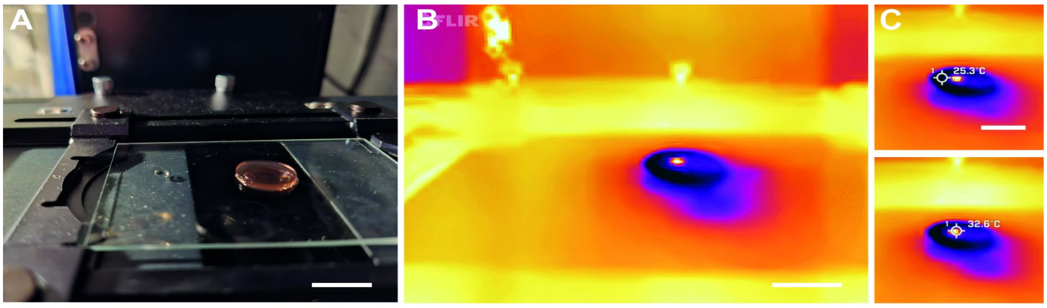

*Figure S4. (A) Unconfined (NIPAM) hydrogel was located under the microscope as a reference. Thermal imaging was done with the FLIR One Edge Pro. (B) The laser, with a power of 153 mW, pulsed on the unconfined NIPAM hydrogel for three cycles (on time 1'000 ms, interval 20'000 ms). The color map iron sheet (Eisenblech) was used for all images. The video of the three cycles running was taken (B) without temperature measurement and with temperature measurement in the (C) center of the laser spot (32.6 °C) as well as the periphery of the NIPAM hydrogel (25.3 °C). The IR images showing here no temperature change in the periphery but a centralized heating that is local around the laser spot. Scale bar 1 cm.*

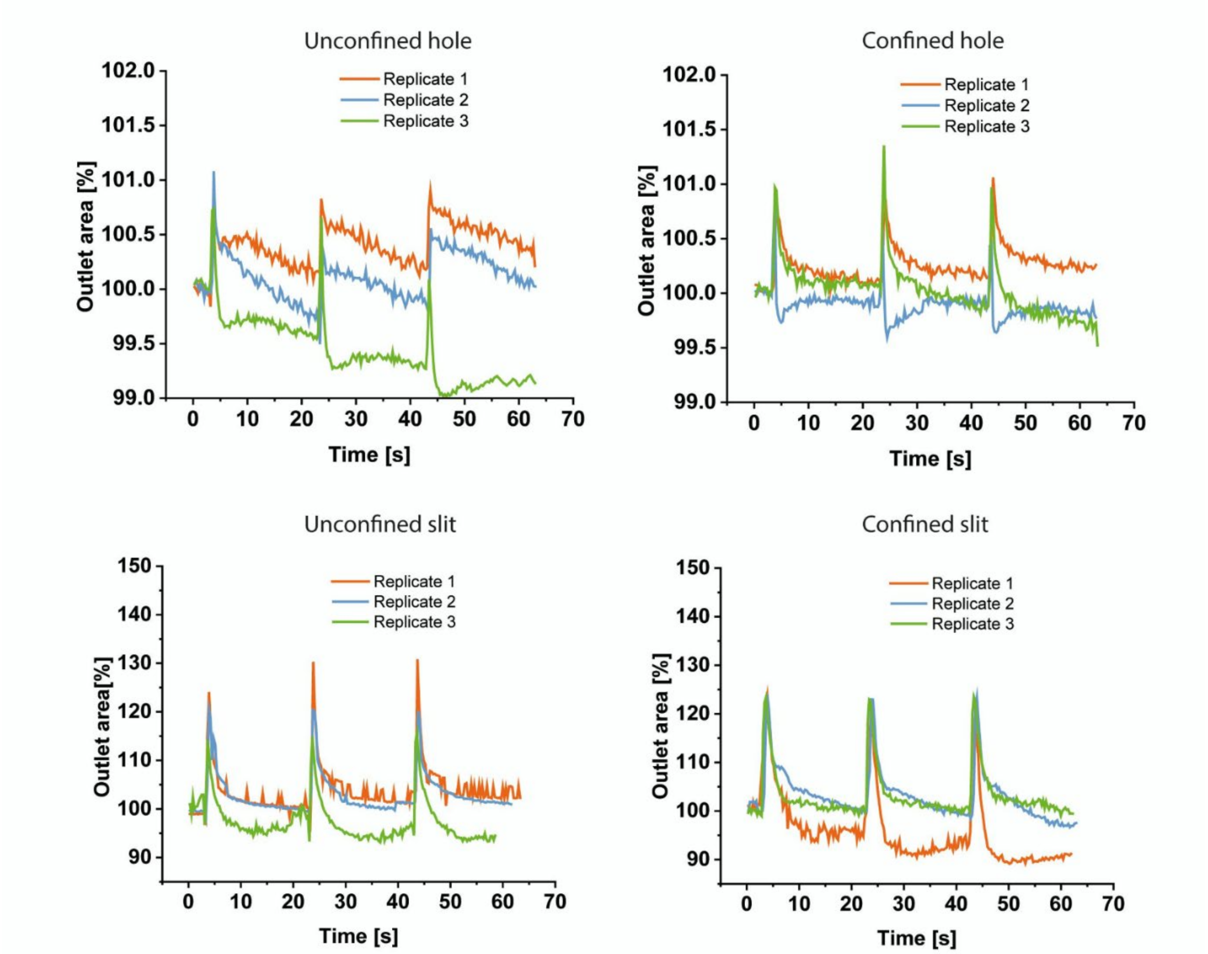

*Figure S5. Replicates of the valves tested as described in Fig 5: unconfined hole, confined hole, unconfined slit and confined slit.*

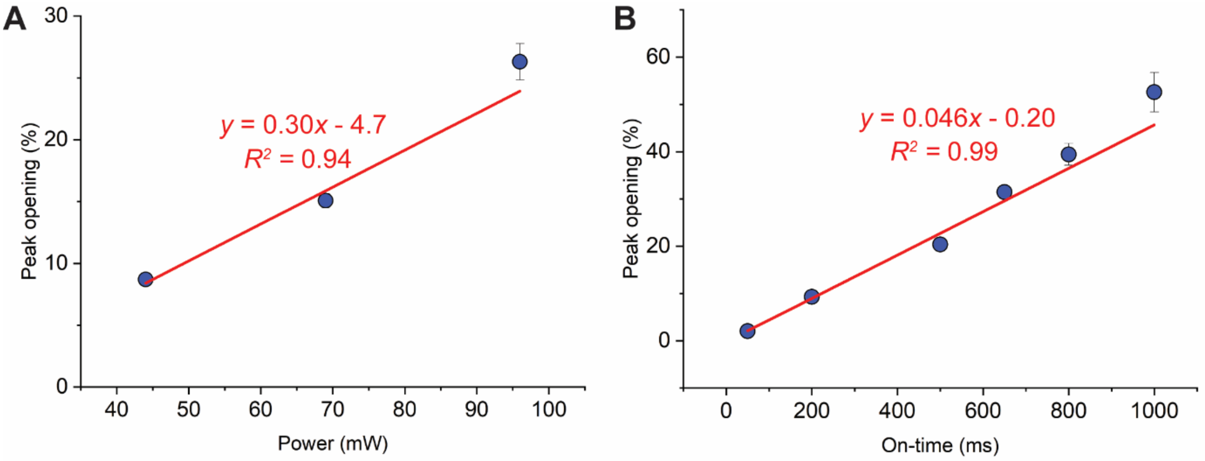

*Figure S6. Effect of changing (A) laser power and (B) on-time on opening of the slit valve. Peak openings (%) are the difference between the maximum size of the valve outlet immediately after a laser pulse and the previous minimum size immediately before the pulse. Each data point in (A) and (B) is an average taken across the 5 or 3 pulses shown in Fig 6A and 6B, respectively. Error bars represent standard errors. Red lines represent linear best fits, with fit parameters and adjusted $R^2$ values shown.*

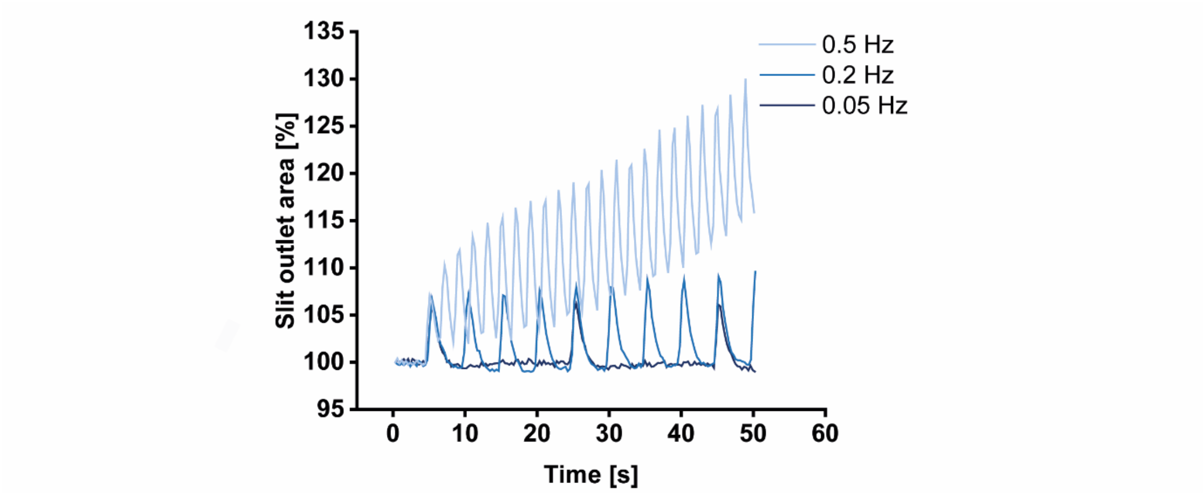

*Figure S7. Effect of changing laser pulse frequency on change in area of the slit outlet. The pulse frequency was varied at constant laser power (96 mW) and pulse duration (500 ms).*

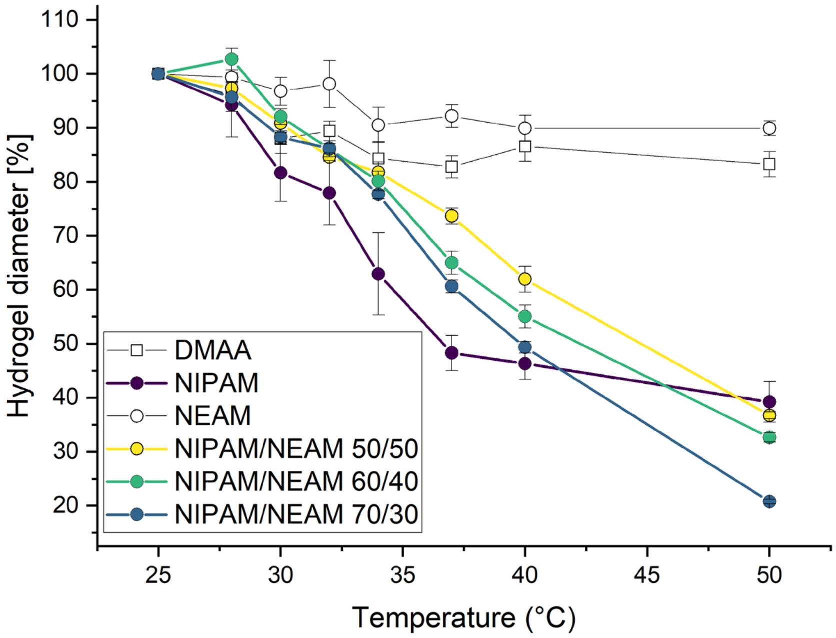

*Figure S8. Diameter of different hydrogels as a function of temperature, measured as a percentage of each gel's diameter at 25 °C. The negative controls of DMAA and NEAM do not change size significantly. Compared to pure NIPAM, copolymers of NIPAM and NEAM have a higher volume phase transition temperature (VPTT).*

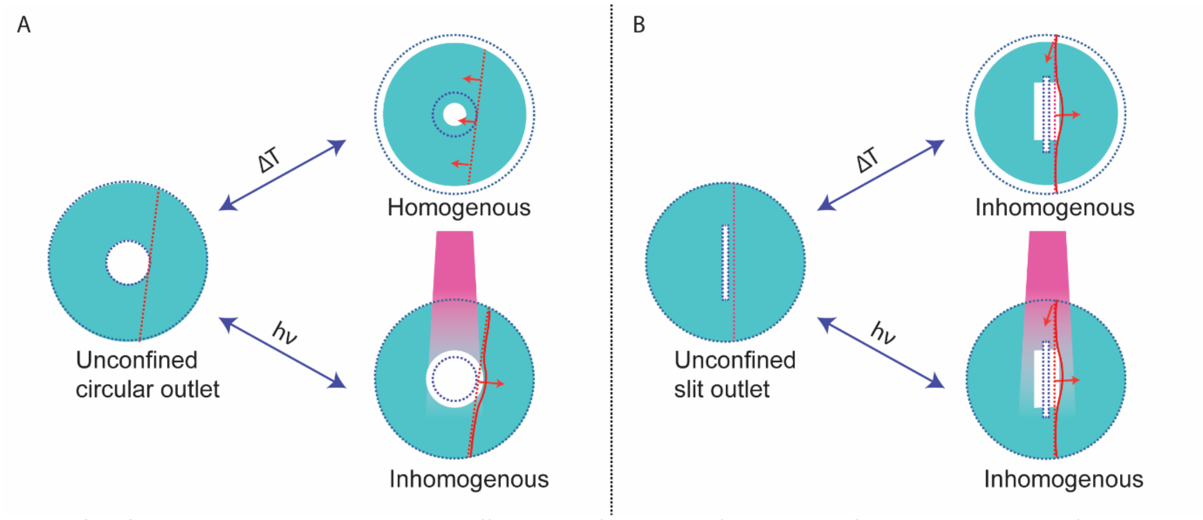

*Figure S9. Changes in temperature or light affect the deformation of two types of outlets in the unconfined hydrogel: (A) circular outlet and (B) slit outlet. These changes result in homogenous or inhomogeneous deformation. Dotted line shows original shape of the outlet.*

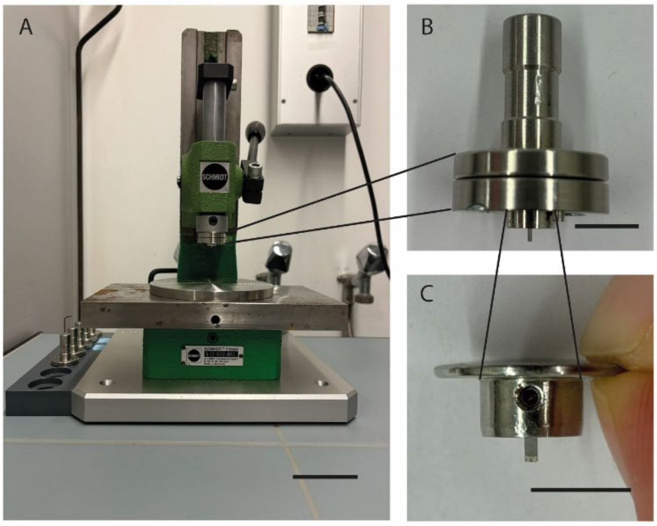

*Figure S10. (A) The mechanical press (scale 5 cm) with (B) the clickable plugin (scale 1 cm) with (C) the beaver blade holder (scale 1 cm), which acts as tongs for the 1 mm beaver blade.*

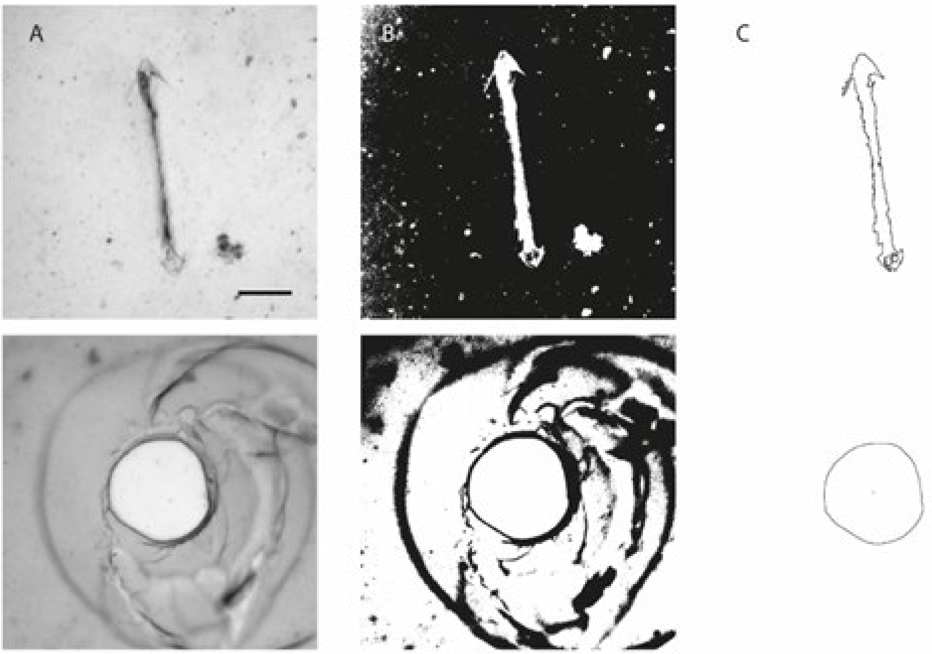

*Figure S11. Evaluation sequence of the outlets containing the (A) original image, (B) auto thresholding using Huang (temperature) or Otsu (light) and (C) drawing of the outlet area. *slit is here only for the light measurement, in the temperature measurement it was measured manually (Fig S12) Scale bar 200 µm.*

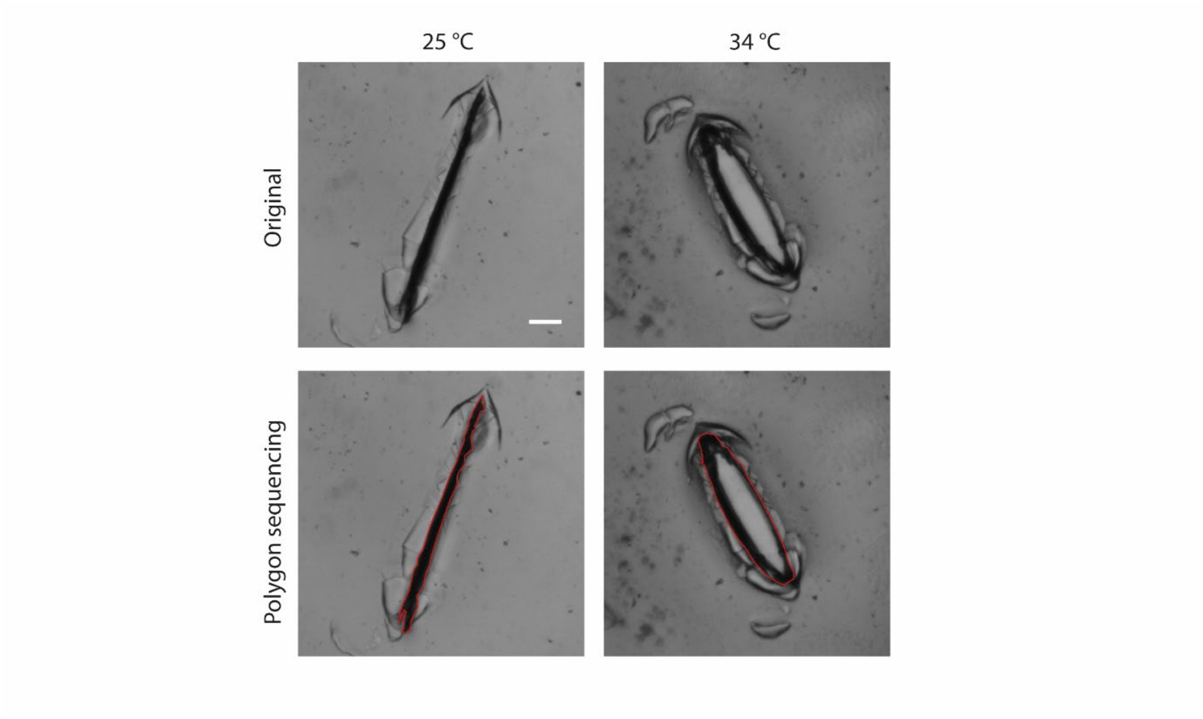

*Figure S12. Example for the slit area measurement in the temperature tests where first the original slit image is shown and then the evaluation of the outlet area using polygon sequencing (by hand) in Fiji where the outermost slit area is being measured.*

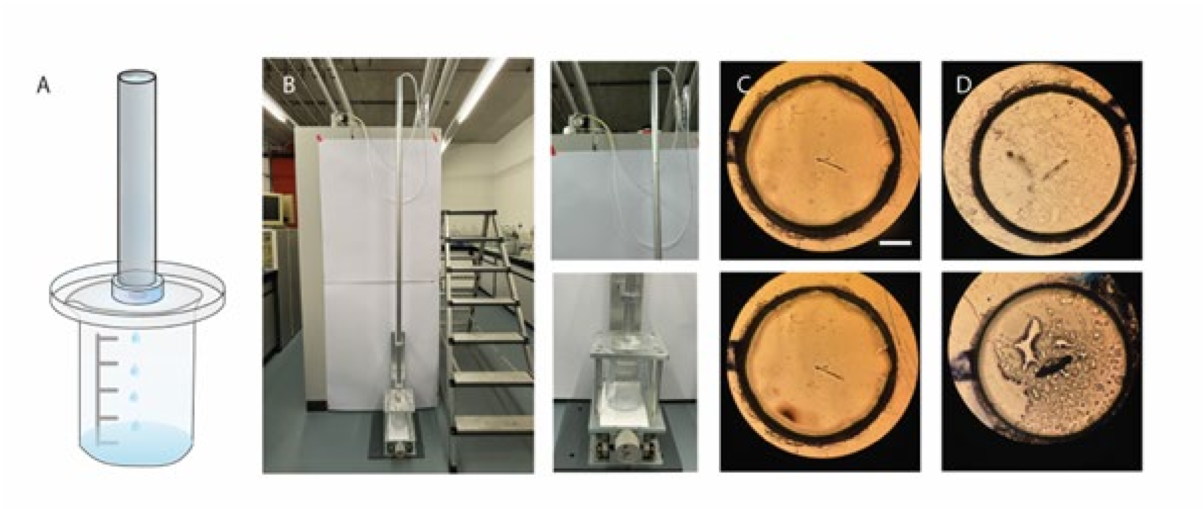

*Figure S13. (A) Schematic of the pressure and flow measurement setup, where the hydrogel is placed on a petri dish with a 4 mm hole and an aluminum rod is tightly aligned with the hydrogel. Between the hydrogel and the aluminum rod, a PDMS ring is placed to prevent the hydrogel from breaking. This setup allows for the simulation of the pressure using hydrostatic pressure. (B) Pressure setup with tubing connected to a pump, enabling the controlled filling of the aluminum rod (I.D.= 2 cm, l =2 m) and a height-adjustable table ensuring a tight seal between hydrogel and rod interface. Position of the slit outlet on the hole of the petri dish before and after (C) pressure measurement and (D) flow measurement. Scale bar 1 mm*

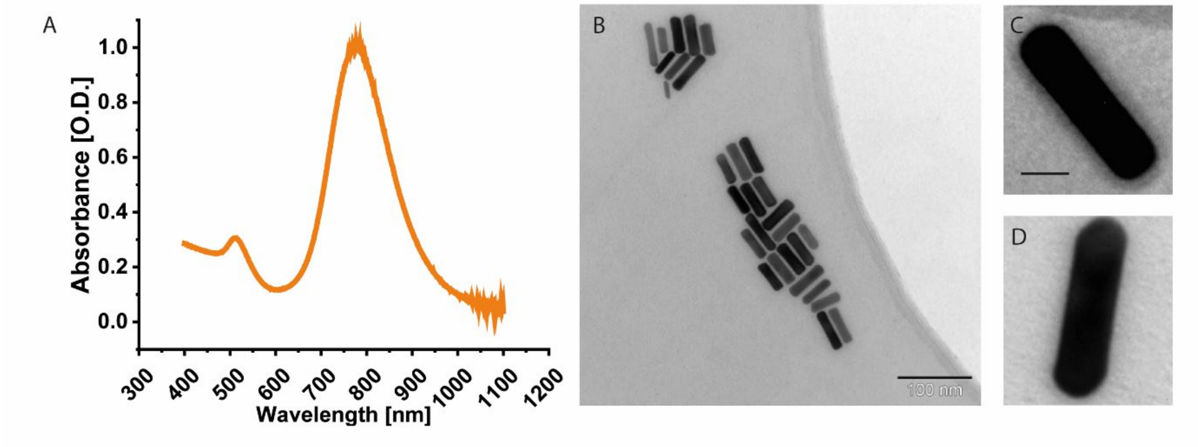

*Figure S14 (A) AuNRs in DMSO with an absorbance peak at 790 nm were produced, which matches the wavelength of the NIR laser at 800 nm. Since no shift of the peak is observed, any aggregation of the NRs is considered negligible. (B) TEM images showing a AuNRs A.R. of 3.7 and the nanorod dimensions of 49 nm in length and 13 nm in width. Scale bar 100 nm. (C) The pegylated (indicated by the dark shadow around the AuNRs) and (D) non-pegylated AuNRs. scale bar 20 nm.*

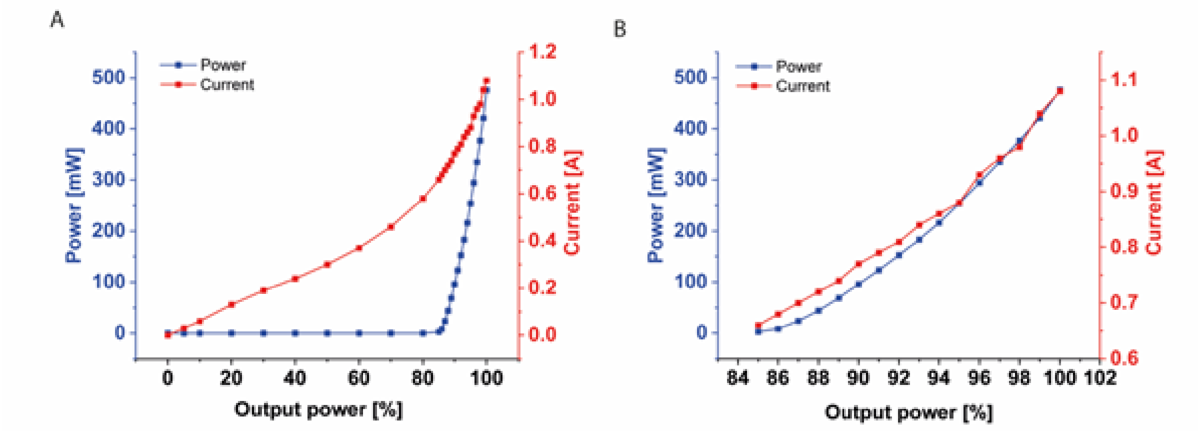

*Figure S15. The laser power output through the adjustable collimator was measured after reducing the laser power at the laser head aperture to 480 mW. A PM2 sensor (2 W, Coherent) in combination with a FieldMax II power meter (Coherent) was used for this measurement. To ensure consistent results, the distance between the collimator and the sensor, as well as the collimator's focus, were kept constant throughout the experiment. (A) Laser power settings below 85% showing that the measured output power remained zero, despite a continuous increase in current. Beyond this threshold, both the output power and current increased continuously. (B) The detailed behavior of the system above 85% laser power, where a nearly linear relationship of power and current is observed. At 100% laser power, the system operated at a current of 1.1 A with a measured output power of 476 mW.*
